# TP53 promotes lineage commitment of human embryonic stem cells through ciliogenesis and sonic hedgehog signaling

**DOI:** 10.1101/2021.07.22.453348

**Authors:** Sushama Sivakumar, Shutao Qi, Ningyan Cheng, Adwait A. Sathe, Mohammed Kanchwala, Ashwani Kumar, Bret M. Evers, Chao Xing, Hongtao Yu

## Abstract

Aneuploidy, defective differentiation, and inactivation of the tumor suppressor *TP53* all occur frequently during tumorigenesis. Here, we probe the potential links among these cancer traits by inactivating *TP53* in human embryonic stem cells (hESCs). *TP53*^−/−^ hESCs exhibit increased proliferation rates, mitotic errors, and low-grade structural aneuploidy; produce poorly differentiated immature teratomas in mice; and fail to differentiate into neural progenitor cells (NPC) *in vitro*. Genome-wide CRISPR screen reveals requirements of ciliogenesis and sonic hedgehog (Shh) pathways for hESC differentiation into NPCs. *TP53* deletion causes abnormal ciliogenesis in neural rosettes. In addition to restraining cell proliferation through *CDKN1A*, TP53 activates the transcription of *BBS9*, which encodes a ciliogenesis regulator required for proper Shh signaling and NPC formation. This developmentally regulated transcriptional program of TP53 promotes ciliogenesis, restrains Shh signaling, and commits hESCs to neural lineages.

## INTRODUCTION

Differentiation status is commonly used to classify the grade of solid tumors, with aggressive high-grade tumors being less differentiated or immature and benign low-grade tumors being more differentiated or mature (Amin et al., 2017). The tumor suppressor gene *TP53* is mutated in more than 50% of all human cancers, and tumors with *TP53* mutations have poor differentiation status (Kastenhuber and Lowe, 2017; Stiewe, 2007). Germline *TP53* mutations cause Li-Fraumeni syndrome (LFS), a hereditary cancer syndrome in humans (Malkin et al., 1990). Patients with LFS develop poorly differentiated breast tumors, and induced pluripotent stem (iPS) cells derived from LFS patient cells exhibit defective osteoblastic differentiation (Lee et al., 2015). Furthermore, *TP53* and its paralogs *TP63* and *TP73* transcriptionally activate the WNT pathway in human and mouse embryonic stem cells (mESCs) to promote mesendodermal specification (Wang et al., 2017). *TP53* negatively regulates self-renewal of neural and myeloid progenitor stem cells by limiting their proliferation and survival (Lin et al., 2005; Zhao et al., 2010). Finally, *TP53* inhibits reprogramming of differentiated cells into iPS cells (Hong et al., 2009; Kawamura et al., 2009; Marion et al., 2009). Collectively, these findings suggest that TP53 antagonizes the stem-cell state and promotes differentiation. By contrast, *Trp53*^−/−^ mice develop normally, although they develop tumors within 6 months of age (Donehower et al., 1992). Moreover, *TP53*-deficient human embryonic stem cells (hESCs) have been reported to form teratomas of all three germ layers when injected into mice (Merkle et al., 2017; Song et al., 2010). These findings argue against a requirement for TP53 in development and stem cell differentiation. Therefore, the functions of TP53 in differentiation remain to be further clarified.

Sonic hedgehog (Shh) signaling regulates tissue patterning during embryogenesis, and its aberrant reactivation in differentiated tissue can lead to tumorigenesis (Liu et al., 2018). In mammals, Shh signaling requires the primary cilium wherein receptors and effectors can be concentrated to high levels and potentiate downstream cellular signaling (Goetz and Anderson, 2010). A well-characterized role for Shh signaling is in neural tube formation and patterning (Goetz and Anderson, 2010). Defects in ciliogenesis perturb Shh signaling, resulting in neural tube defects. hESCs have been reported to possess primary cilia, which are enriched for Shh pathway components (Kiprilov et al., 2008).

Aneuploidy is a common trait of tumor cells, with about 90% of human solid tumors exhibiting aneuploidy (Ben-David and Amon, 2020). High-degree aneuploidy is linked to aggressive, metastatic tumors with poor prognosis. Studies using yeast, mouse embryonic fibroblasts and human Down syndrome fibroblasts have clearly indicated that the gain of extra chromosomes alters gene dosage and inhibits cell proliferation (Ben-David and Amon, 2020). On the other hand, aneuploid tumor cell lines proliferate with high fitness in culture, suggesting that cancer cells might have lost aneuploidy-suppressing mechanisms during tumor evolution *in vivo*. TP53 mutations are associated with high-degree aneuploidy in human cancers (Pfister et al., 2018), suggesting a role of TP53 in suppressing aneuploidy. Yet, inactivation of *TP53* in cultured diploid or near-diploid cells fail to induce gross aneuploidy (Bunz et al., 2002). These paradoxical findings suggest the existence of selective pressures *in vivo* that allow aneuploid cells to gain higher fitness over their euploid counterparts.

Embryonic stem cells (ESCs) are prone to develop spontaneous aneuploidy and tend to acquire *TP53* mutations in culture (Merkle et al., 2017; Zhang et al., 2016). Trisomy of single chromosomes in mESCs has been reported to cause increased proliferation rates, decreased differentiation capacity *in vitro* and the formation of immature teratomas *in vivo* (Zhang et al., 2016). Human teratomas are germ cells tumors that consist of several different tissue types. Interestingly, virtually all testicular germ cell tumors exhibit aneuploidy (Shen et al., 2018). These findings suggest that aneuploidy might be more tolerated in ESCs.

In an effort to understand the origins of aneuploidy, we sought to use teratomas grown from hESCs in mice as an artificial *in vivo* model to evolve aneuploidy. We inactivated *TP53* in hESCs with CRISPR-Cas9 and examined the karyotype, transcriptome, and differentiation of *TP53*-deficient hESCs *in vitro* and using the teratoma assay. Our results indicate that *TP53* loss is insufficient to produce gross whole-chromosome aneuploidy in hESCs or during teratoma formation. Unexpectedly, we discovered that *TP53*-deficient hESCs formed poorly differentiated, immature teratomas and failed to differentiate into functional NPCs *in vitro*. Genome-wide CRISPR-Cas9 screening, gene expression profiling, and ChIP-seq experiments further link TP53 to ciliogenesis and Shh signaling during NPC formation. Therefore, TP53 has an underappreciated role in lineage commitment of hESCs.

## RESULTS

### *TP53* inactivation in hESCs does not cause prevalent whole-chromosome aneuploidy

We inactivated *TP53* in both H1 and H9 hESCs using CRISPR-Cas9, with either plasmid transfection or lentiviral infection and multiple sgRNAs that targeted the transactivation domain (TAD) and the DNA-binding domain (DBD) of TP53 (Figures 1A and S1A). Among the five *TP53*^−/−^ hESC lines generated, H9 *TP53*^−/−^ clones 1 and 2 (C1 and C2) were derived from single clones, whereas the others were mixed cell populations to avoid clonal bias. As expected, the full-length (FL) TP53 protein was absent in all *TP53*^−/−^ hESC lines (Figure 1B). H1 *TP53*^−/−^, H9 *TP53*^−/−^ C1, and H9 *TP53*^−/−^ lines expressed truncated forms of TP53, namely ΔN40 or ΔN133 TP53 (Aoubala et al., 2011; Courtois et al., 2002). These truncated forms of TP53 are functionally deficient, as they lack an intact TAD (Figure 1A). The *TP53*^−/−^ hESCs maintained the expression of pluripotency markers NANOG, SOX2, and OCT3/4, based on flow cytometry and immunofluorescence (Figures 1C and S1B).

**Figure 1.**
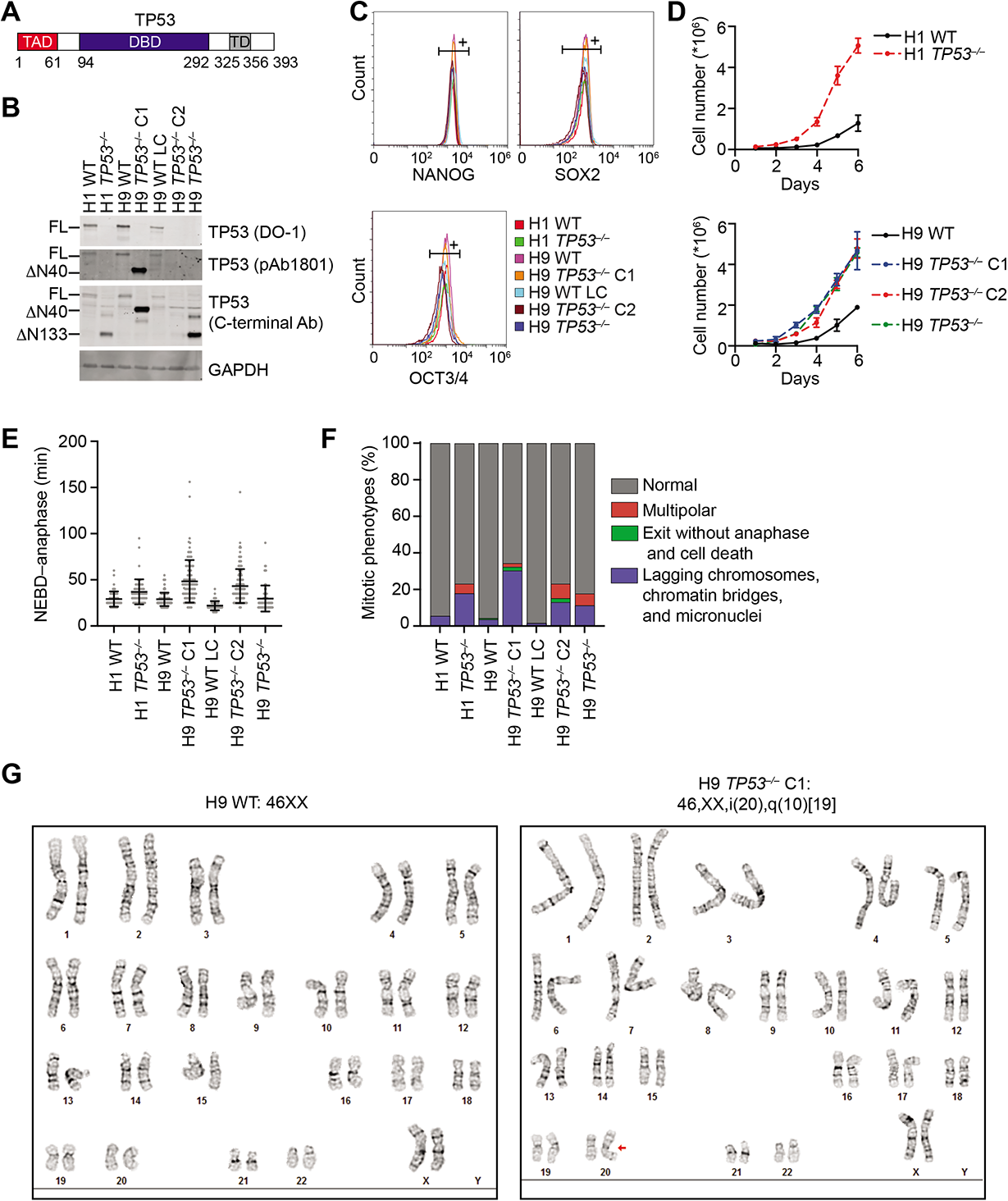
*TP53^−/−^* hESCs exhibit mitotic errors but no whole-chromosome aneuploidy. (A) Schematic representation of the domains of TP53. TAD, transactivation domain; DBD, DNA binding domain; TD, tetramerization domain. (B) Western blots of total cell lysates of *TP53^−/−^* hESC clones and pools. The DO-1 antibody detects full-length (FL) TP53. pAb1801 and the TP53 C-terminal antibody detect the N-terminal truncation of TP53 (ΔN40 and ΔN133). (C) Flow cytometry analysis of wild type (WT) and *TP53^−/−^* hESCs stained with pluripotency markers. (D) Cell proliferation assays of WT and *TP53^−/−^* hESCs. Mean ± s.d., n=3 independent experiments. (E) Quantification of the mitotic durations from nuclear envelope breakdown (NEBD) to anaphase onset by live-cell imaging of WT and *TP53^−/−^* hESCs. Each dot represents one cell. Mean ± s.d.; n=50-100 cells. (F) Quantification of the indicated mitotic phenotypes of WT and *TP53^−/−^* hESCs as revealed by live-cell imaging. (G) Karyotypes of H9 WT and *TP53^−/−^* C1 hESCs by Giemsa banding. Red arrow indicates isochromosome 20q.

The *TP53*^−/−^ hESCs proliferated faster than wild-type (WT) hESCs (Figure 1D). *TP53*^−/−^ hESCs had larger nuclear and cell sizes, but their ratios were not greatly altered (Figures S1C-E). Consistent with a previous report (Drost et al., 2015), *TP53* inactivation caused delays in mitotic progression and increased incidence of lagging chromosomes, chromatin bridges, micronuclei, multipolar spindles, and mitotic slippage (Figures 1E and 1F). Despite the larger nuclear size and the increased mitotic errors, *TP53*^−/−^ hESCs had the normal diploid karyotype and did not display prevalent polyploidy and whole-chromosome aneuploidy, as revealed by karyotype analysis (Table S1). Culturing *TP53*^−/−^ hESCs for an additional 21 passages did not increase the extent of aneuploidy.

Some *TP53*^−/−^ hESCs exhibited arm-level copy number variations (CNVs) in certain chromosomes. For example, the H9 *TP53*^−/−^ C1 line had an isochromosome i20q10 abnormality, which was caused by deletion of 20p and duplication of 20q (Figure 1G). This abnormality was also present in 10% of the pooled population of H9 *TP53*^−/−^ hESCs. The i20q10 structural aberration is commonly found in long-term cultures of hESCs (Taapken et al., 2011). *TP53* loss likely accelerated the emergence of this abnormality in *TP53*^−/−^ hESCs.

*TP53*^−/−^ hESCs had increased mitotic defects but did not exhibit aneuploidy, suggesting that aneuploidy might not be tolerated in these cells. To further test this notion, we treated WT and *TP53*^−/−^ hESCs with reversine, a chemical inhibitor of the spindle checkpoint kinase MPS1 (Santaguida et al., 2010). Inhibition of MPS1 is known to inactivate the spindle checkpoint and cause premature chromosome segregation prior to the establishment of proper kinetochore-microtubule attachment. WT hESCs did not survive reversine treatment, whereas *TP53*^−/−^ hESCs survived and could be propagated. Karyotype analysis showed that reversine-treated *TP53*^−/−^ hESCs did not exhibit whole-chromosome aneuploidy (Table S1). Interestingly, reversine treatment increased the incidence of the i20q10 structural aberration in these cells. Taken together, our results indicate that TP53 loss is insufficient to induce whole-chromosome aneuploidy in hESCs, but permits the occurrence of arm-level CNVs.

### *TP53^-/-^* hESCs form immature teratomas

When injected into immunodeficient mice, hESCs develop into teratomas that contain cells from all three germ layers (ectoderm, mesoderm, and endoderm) (Solter, 2006). The formation of teratomas *in vivo* is an important test for pluripotency in stem cells. To test the pluripotency of *TP53*^−/−^ hESCs, we injected WT and *TP53*^−/−^ hESCs subcutaneously into immunodeficient NOD-SCID mice. Macroscopically, the WT teratomas contained multiple fluid-filled cysts, while the *TP53*^−/−^ teratomas were solid tumors (Figure S2A). Histological analysis revealed that the WT teratomas were composed of large cysts lined with mature, well-differentiated cells as well as other mature tissues, belonging to all three germ layers (Figure S2B). By contrast, the *TP53*^−/−^ teratomas were predominately composed of sheets of poorly differentiated cells with high nuclear to cytoplasmic (NC) ratio. Quantification confirmed that WT teratomas indeed consisted mostly of mature elements whereas *TP53*^−/−^ teratomas contained high percentages of poorly differentiated, immature elements (Figures 2A and 2B). Large neural rosette-like structures formed by poorly differentiated neural cells were frequently observed in *TP53*^−/−^ teratomas. WT teratomas contained many ciliated cells and few mitotic cells (Figures S3A and S3B). The *TP53*^−/−^ teratomas, however, lacked ciliated cells and exhibited higher mitotic indices. Compared to WT teratomas, expression of the mature element marker GFAP was decreased in *TP53*^−/−^ teratomas while expression of SALL4 and Glypican-3, which are markers for immature elements, was increased (Figure S3C). Taken together, our results clearly show that, contrary to a previous report (Song et al., 2010), deletion of *TP53* impairs differentiation of hESCs in teratomas *in vivo*.

**Figure 2.**
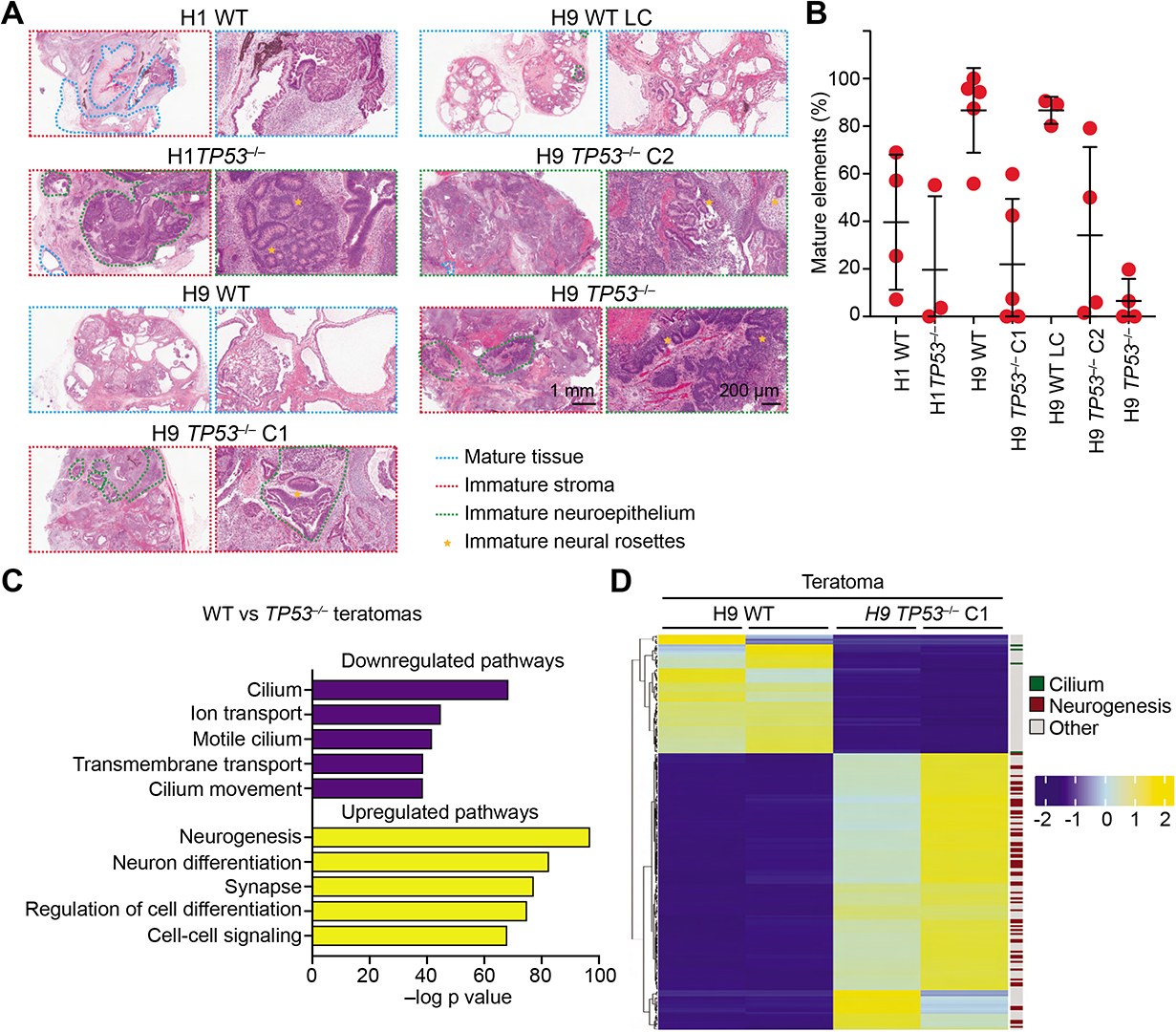
*TP53^−/−^* hESCs form immature teratomas with deficient neurogenesis and ciliogenesis. (A) Hematoxylin and eosin (H&E) staining of WT and *TP53^−/−^* teratoma sections. Mature elements, immature stroma, immature neuroepithelium, and immature neural rosettes are marked by dashed lines of the indicated color. Immature neural rosettes are marked by stars. (B) Quantification of the percentages of mature and immature elements in WT and *TP53^−/−^* teratoma sections. Mean ± s.d., n=3-5. (C) Gene set enrichment analysis (GSEA) of DEGs between WT and *TP53^−/−^* teratomas. (D) Heatmap of top 200 differentially expressed genes (DEGs) identified by RNA-seq analysis of WT and *TP53^−/−^* teratomas. DEGs involved in the cilium and neurogenesis pathways are indicated.

### *TP53*^−/−^ teratomas do not exhibit gross aneuploidy

Loss of TP53 alone is insufficient to produce aneuploidy in hESCs cultured *in vitro*, a finding that is in general agreement with the proliferation-suppressive function of aneuploidy (Ben-David and Amon, 2020). We next tested whether the highly immature and poorly differentiated *TP53*^−/−^ teratomas might foster aneuploidy *in vivo* by detecting CNVs with whole genome sequencing (WGS). As expected, WT hESCs and teratomas were diploid (Figures S4A and S4B). As described above, based on karyotyping, H9 *TP53*^−/−^ C1 cells harbored the i20q10 abnormality that resulted in trisomy for 20q and monosomy for 20p. WGS analysis of H9 *TP53*^−/−^ C1 cells reliably detected the presence of these CNVs. Likewise, as revealed by karyotyping, about 60% of H9 *TP53*^−/−^ C2 cells harbored deletion of a large segment of chromosome 18q. This CNV was detected by WGS analysis of H9 *TP53*^−/−^ C2 cells, which also detected the duplication of a small region of chromosome 12p, an abnormality not detected by karyotyping. The CNVs observed in these cultured *TP53*^−/−^ hESCs prior to their injection into mice were also detected in teratomas grown from these cells. Surprisingly, despite their highly immature nature, these *TP53*^−/−^ teratomas did not exhibit additional whole-chromosome or even arm-level aneuploidy (Figures S4A, S4B, and Table S1).

Interestingly, about 10% of the cells in the H9 *TP53*^−/−^ pool exhibited the i20q10 abnormality while about 50% of the cells in the teratoma formed by these cells contained this aberration (Figures S4A, S4B, and Table S1). Thus, this arm-level aneuploidy was enriched during teratoma formation, suggesting that cells with this chromosome abnormality might have a proliferation advantage over diploid cells. This observation also indicated that genetic alterations conferring proliferation advantage could be amplified and fixed in the cell population during the relatively short duration of teratoma formation. Therefore, the fact that *TP53*^−/−^ teratomas lack gross aneuploidy suggests that TP53 loss is insufficient to tolerate aneuploidy during teratoma formation.

### *TP53*^−/−^ hESCs are deficient in differentiation into neural lineages

To characterize the differentiation defects of *TP53*^−/−^ teratomas, we compared the gene expression profiles of WT and H9 *TP53*^−/−^ teratomas with RNA-seq. About 3,000 differentially expressed genes (DEGs) were identified between WT and *TP53*^−/−^ teratomas, indicative of global gene expression changes. Tissue specific enrichment analysis (TSEA) showed that the top 200 DEGs were enriched for genes specifically expressed in the brain (Figure S4C). Pathway analysis of the DEGs revealed that the cilium pathway was downregulated in *TP53*^−/−^ teratomas whereas neurogenesis and neuron differentiation were among the top upregulated pathways (Figure 2C). Heatmap of the top 200 DEGs revealed that multiple ciliary genes and numerous neurogenesis genes were down-and up-regulated, respectively, in *TP53*^−/−^ teratomas (Figure 2D). The gene expression changes are thus consistent with the histological observation that *TP53*^−/−^ teratomas lacked cilia and were enriched for immature neural elements.

To probe the functions of TP53 in neural differentiation, we performed an *in vitro* differentiation assay of hESCs using the 3D embryoid body (EB) protocol (Figure S5A). In this assay, hESCs are first plated in microwells to form EBs. The EBs are then plated in monolayers to form neural rosettes (NRs), a radial arrangement of cells representative of the neural tube stage in embryonic development (Deglincerti et al., 2016; Elkabetz et al., 2008). NRs are then isolated and further induced to produce neural progenitor cells (NPCs) that actively proliferate. We analyzed the ability of WT and *TP53*^−/−^ hESCs to form EB, NR and NPCs during differentiation. Both WT and *TP53*^−/−^ hESCs formed EBs efficiently, with the *TP53*^−/−^ EBs being slightly smaller than WT controls (Figures S5B and S5C).

WT EBs expectedly produced well-organized NRs, as evidenced by the radial arrangement of the tight junction protein ZO-1, which redistributed to the apical side of NRs during differentiation (Elkabetz et al., 2008) (Figure 3A). Another feature of mature NRs is the generation and organization of primary cilia in the apical region (He et al., 2014). As evidenced by staining with centriole and ciliary markers and acetylated tubulin, WT NRs contained radial arrays of cilia with their tips pointing towards the apical lumen (Figures 3B and 3C). By contrast, *TP53*^−/−^ EBs were severely impaired in NR organization. The total differentiated areas of *TP53*^−/−^ NRs were greatly decreased (Figures S5D and S5E). In *TP53*^−/−^ NRs, expression of the NR marker PAX6 was deficient (Figure S5F). *TP53*^−/−^ NRs were disorganized, as evidenced by the incomplete ZO-1 redistribution to the apical region and disordered arrays of acetylated tubulin and cilia (Figures 3A-3C). Furthermore, compared to that of WT NR, the number of radial arrangements in each *TP53*^−/−^ NR was also reduced (Figure 3D).

**Figure 3.**
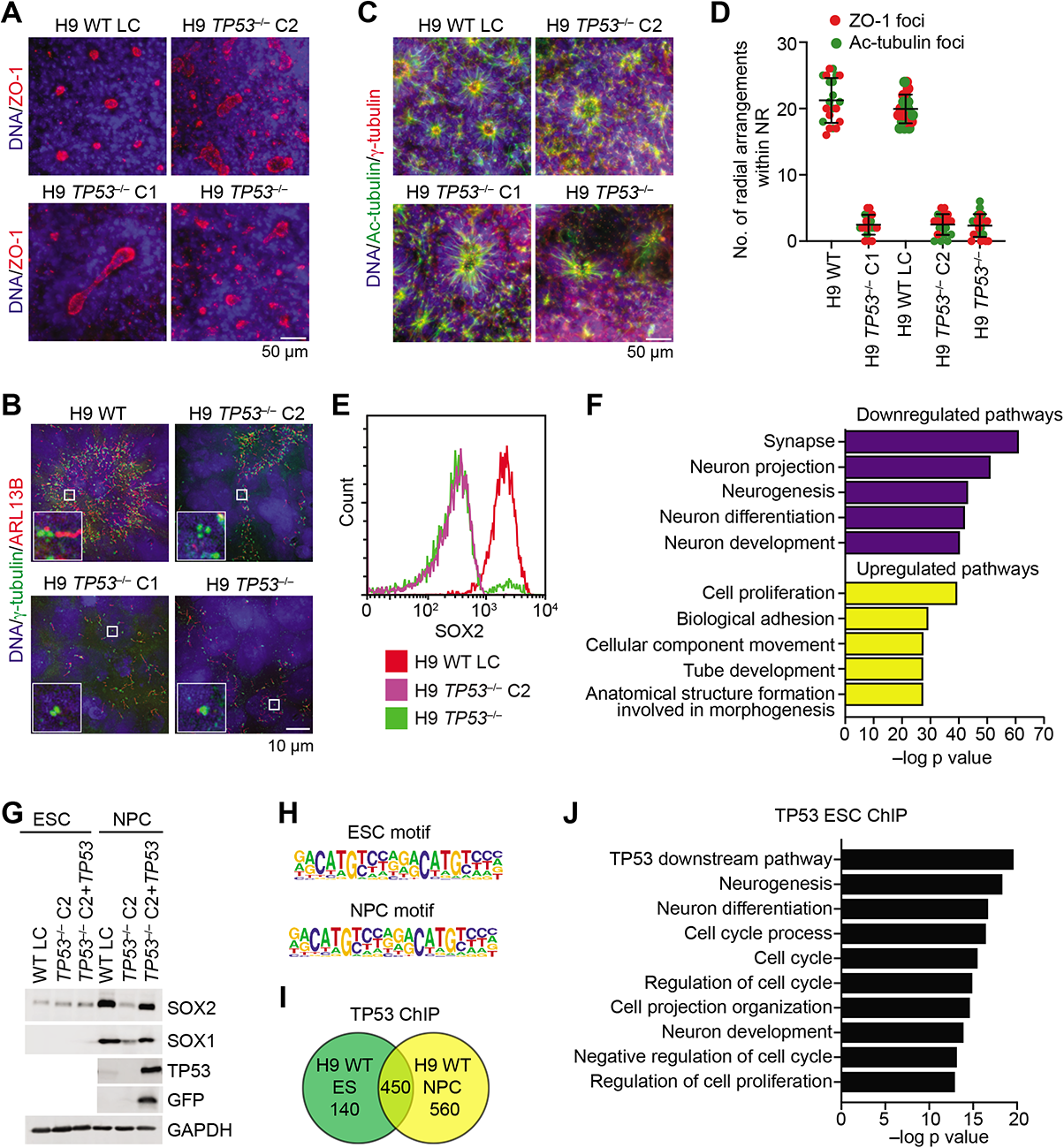
*TP53^−/−^* hESCs are deficient in differentiation into neural progenitor cells (NPCs) *in vitro*. (A) Immunofluorescence (IF) images of WT and *TP53^−/−^* neural rosettes (NRs) at day 11 stained with DAPI for DNA (blue) and the ZO-1 antibody (red). (B) IF images of WT and *TP53^−/−^* NRs at day 11 stained with DAPI for DNA (blue) and antibodies against the cilium protein ARL13B (red) and γ-tubulin (green). (C) IF images of WT and *TP53^−/−^* NRs at day 11 stained with DAPI for DNA (blue) and antibodies against acetylated tubulin (Ac-tubulin; green) and γ-tubulin (red). (D) Quantification of the number of radial arrangements (as defined by ZO-1 and Ac-tubulin foci) within each WT or *TP53^−/−^* NR. Each dot represents one NR. Mean ± s.d.; n=10. (E) Flow cytometry analysis of SOX2 expression in WT and *TP53^−/−^* NPCs. (F) Gene set enrichment analysis (GSEA) of the 589 common DEGs between WT and *TP53^−/−^* NPCs. (G) Western blots of total lysates of WT and *TP53^−/−^* ESCs and NPCs and *TP53^−/−^* cells expressing an exogenous *TP53* transgene. (H) *De novo* motif analysis of TP53 ChIP-seq peaks in ESCs and NPCs. (I) Venn diagram showing the number and overlap of TP53 ChIP-seq peaks in WT ESCs and NPCs. (J) GSEA of genes occupied by TP53 in ESCs.

The WT NRs efficiently differentiated into NPCs that expressed NPC markers SOX1, SOX2 and Nestin based on flow cytometry, immunostaining, and Western blotting (Figures 3E and S5G-S5J). The differentiation of *TP53*^−/−^ NRs into NPCs was defective, as *TP53*^−/−^ NPCs displayed greatly reduced expression of SOX1, SOX2 and Nestin. The NPC-like cells derived from *TP53*^−/−^ NRs were still capable of proliferation and expressed similar levels of the proliferation marker KI67 (Figures S6A and S6B). *TP53*^−/−^ NPCs were functionally defective, as they failed to differentiate into astrocytes or neurons, as evidenced by decreased expression of both the astrocyte marker GFAP and neuronal markers MAP2 and β-III-Tubulin (Figure S6C, S6D).

Although TP53 loss did not cause gross aneuploidy in ESCs, it did create genomic instability, including arm-level aneuploidy, which was different in each *TP53^−/−^* hESC line. Indeed, karyotype analysis revealed that 20-25% of NPCs derived from two different *TP53^−/−^* hESC lines retained the corresponding original structural aneuploidy in the ESC state (Figure S6E and Table S1).

To ascertain the extent of differentiation defects of *TP53*^−/−^ NPCs, we compared the gene expression profiles of NPCs derived from three H9 *TP53*^−/−^ hESC lines with their respective WT controls. Pairwise comparisons revealed 1660 DEGs between NPCs derived from H9 WT and *TP53*^−/−^ C1, 3113 DEGs between H9 WT and *TP53*^−/−^ C2, and 2131 DEGs between H9 LC and the *TP53^−/−^* pool (Figure S7A). We reasoned that common DEGs found in all three pairwise comparisons might be less likely influenced by genomic instability caused by TP53 loss and capture the genuine differentiation defects in *TP53^−/−^* NPCs. Only 589 DEGs were common among all three pairwise comparisons.

The heatmap of expression levels of these 589 common genes showed that multiple genes involved in neuron differentiation were downregulated whereas genes involved in cell proliferation were upregulated (Figure S7B). Pathway analysis further confirmed that the top downregulated pathways were integral to neuronal functions and differentiation (Figure 3F). The top upregulated pathways included cell proliferation, biological adhesion, cellular component movement and tube development. Thus, *TP53*^−/−^ NPCs are deficient in the expression of genes required for neuronal differentiation.

### TP53 occupies neurogenesis genes in hESCs

Multiple *TP53^−/−^* hESCs with distinct chromosome abnormalities and genetic mutations exhibited similar differentiation defects in the teratoma and *in vitro* differentiation assays. Expression of the WT *TP53* transgene in H9 *TP53^−/−^* C2 hESCs restored SOX1 and SOX2 expression in the NPC state (Figures 3G, S5G, and S5H). Therefore, the genomic instability unleashed by TP53 loss is not solely responsible for the differentiation defects. As a transcription factor, TP53 might directly or indirectly promote the transcription of cell fate determination genes, including *SOX1* and *SOX2*.

To identify direct transcription targets of TP53, we performed chromatin immunoprecipitation followed by next-generation sequencing (ChIP-seq) analysis in WT hESCs and NPCs. *De novo* motif search identified the TP53-binding consensus in the enriched peaks in both hESCs and NPCs (Figure 3H), confirming the validity of the ChIP-seq data. About 76% of the chromosome regions occupied by TP53 in the stem cell state remained occupied by TP53 in the NPC state (Figures 3I and S7C). Pathway analysis of genes occupied by TP53 in hESCs unexpectedly reveals the enrichment of genes in the neurogenesis and neuron differentiation pathways (Figure 3J), aside from the expected enrichment of genes in the canonical TP53 pathways. This surprising finding suggests that TP53 already occupies genes required for neuronal differentiation in the stem cell state.

RNA-seq analysis showed that 106 DEGs were common among pairwise comparisons between three different *TP53^−/−^* hESCs and their corresponding WT controls (Figure S7D). The expression heatmap of the common 106 DEGs showed that multiple genes involved in the TP53 pathway were downregulated whereas genes involved in cell proliferation were upregulated (Figure S7E). Pathway analysis of these DEGs expectedly identified the canonical TP53 pathways, but did not show enrichment of neurogenesis genes (Figure S7F). Despite being occupied by TP53 in hESCs, several known regulators of neuronal functions, including *SYT1* and *SLITRK1*, were not differentially expressed between WT and *TP53^−/−^* hESCs. By contrast, 54% of the TP53-occupied genes in NPCs were differentially expressed between WT and *TP53^−/−^* NPCs. Thus, TP53 already occupies neuronal genes at the stem cell state and is poised to activate their expression during NPC differentiation.

### Genome-wide screen reveals requirement for ciliogenesis and Shh signaling during NPC differentiation

To functionally link TP53 target genes to differentiation, we performed a genome-wide CRISPR-Cas9 screen to identify genes required for NPC differentiation. RNA-seq analysis revealed that the well-established NPC marker *SOX1* was indeed the top DEG between WT hESCs and NPCs (Figure S8A). We thus decided to use SOX1 as the surrogate marker for NPCs (Figure 4A). WT hESCs were infected with the lentiviral human GeCKOv2 library and induced to undergo NPC differentiation by the monolayer protocol (Chambers et al., 2009). We collected the 1% of cells with lowest SOX1 expression from the differentiated cell population using flow cytometry. The sgRNAs enriched in these SOX1^−^ cells were thus expected to target genes required for NPC differentiation. *SOX1* was a top hit from the screen, confirming its validity (Figure 4B). Other top hits included genes involved in ciliogenesis and negative regulators of Shh signaling (Figures 4B and 4C). Our screen was not saturated, as *TP53* was not identified in the screen, despite the fact that *TP53^−/−^* hESCs, when subjected to this monolayer protocol, were deficient in NPC differentiation and SOX1 expression.

**Figure 4.**
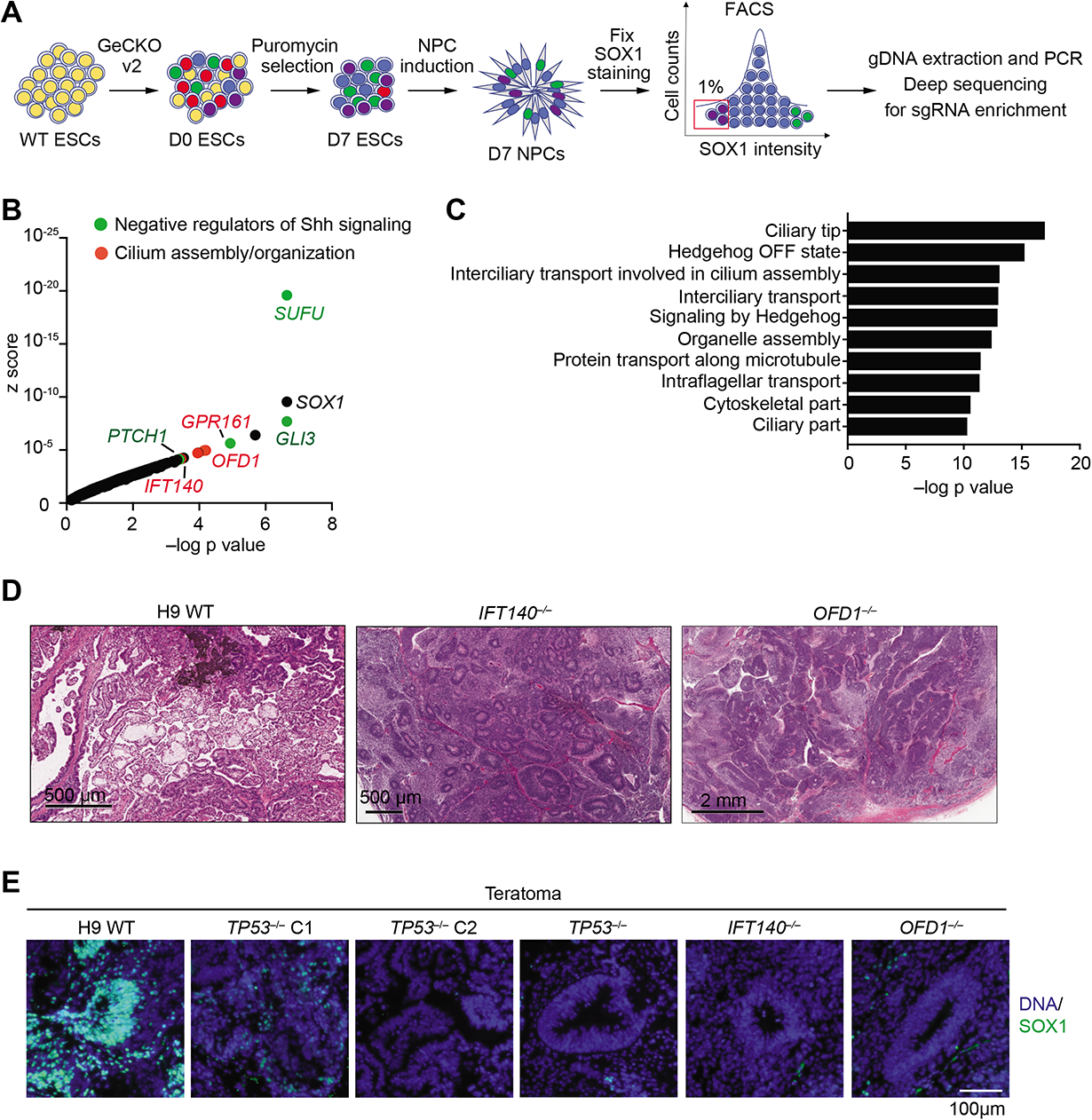
Genome-wide CRISPR screen identifies requirement for ciliogenesis and downregulation of sonic hedgehog (Shh) signaling during NPC differentiation. (A) Flow chart of the CRISPR-Cas9 screen. (B) Graphical display of CRISPR-Cas9 screen hits ranked by the z score. Top 20 hits with known functions in Shh signaling and ciliogenesis are depicted in green and red dots, respectively. (C) Gene Set Enrichment Analysis (GSEA) of top 500 hits in the CRISPR screen. (D) Hematoxylin and eosin (H&E) staining of WT, *IFT140^−/−^* and *OFD1^−/−^* teratoma sections at high magnification. Four teratomas in each group were analyzed with representative images shown. (E) Immunohistochemistry (IHC) staining of the indicated teratoma sections with DAPI for DNA and the anti-SOX1 antibody.

The ciliary genes *IFT140* and *OFD1* were among the top hits in our screen. IFT140 is a core subunit of the IFT-A subcomplex, which is required for retrograde transport within the primary cilium (Picariello et al., 2019). OFD1 is required for ciliogenesis and neuronal patterning in mice and in mouse stem cells (Ferrante et al., 2006; Hunkapiller et al., 2011). To study the functions of ciliogenesis and Shh signaling during NPC differentiation, we generated hESCs with *IFT140* and *OFD1* individually deleted (Figure S8B). About 30% of WT hESCs possessed primary cilia (Figures S8C and S8D). Deletion of *IFT140* or *OFD1* abolished ciliogenesis in hESCs. Strikingly, when injected into mice, *IFT140^−/−^* and *OFD1^−/−^* hESCs formed highly immature teratomas with numerous disorganized, less compact NRs (Figures 4D and S8E). Similar to *TP53^−/−^* teratomas, *IFT140^−/−^* and *OFD1^−/−^* teratomas exhibited greatly reduced SOX1 expression (Figures 4E), confirming our screen results. Ciliogenesis in NRs of *IFT140^−/−^* and *OFD1^−/−^* teratomas was severely compromised (Figure S8F). Similar defects in NR organization, ciliogenesis, and SOX1 expression were also observed for *IFT140^−/−^* and *OFD1^−/−^* hESCs during *in vitro* differentiation (Figures 5A-5D). Taken together, our results suggest that ciliogenesis promotes the formation of tightly compacted NRs, which are critical for differentiation towards neuronal lineages.

**Figure 5.**
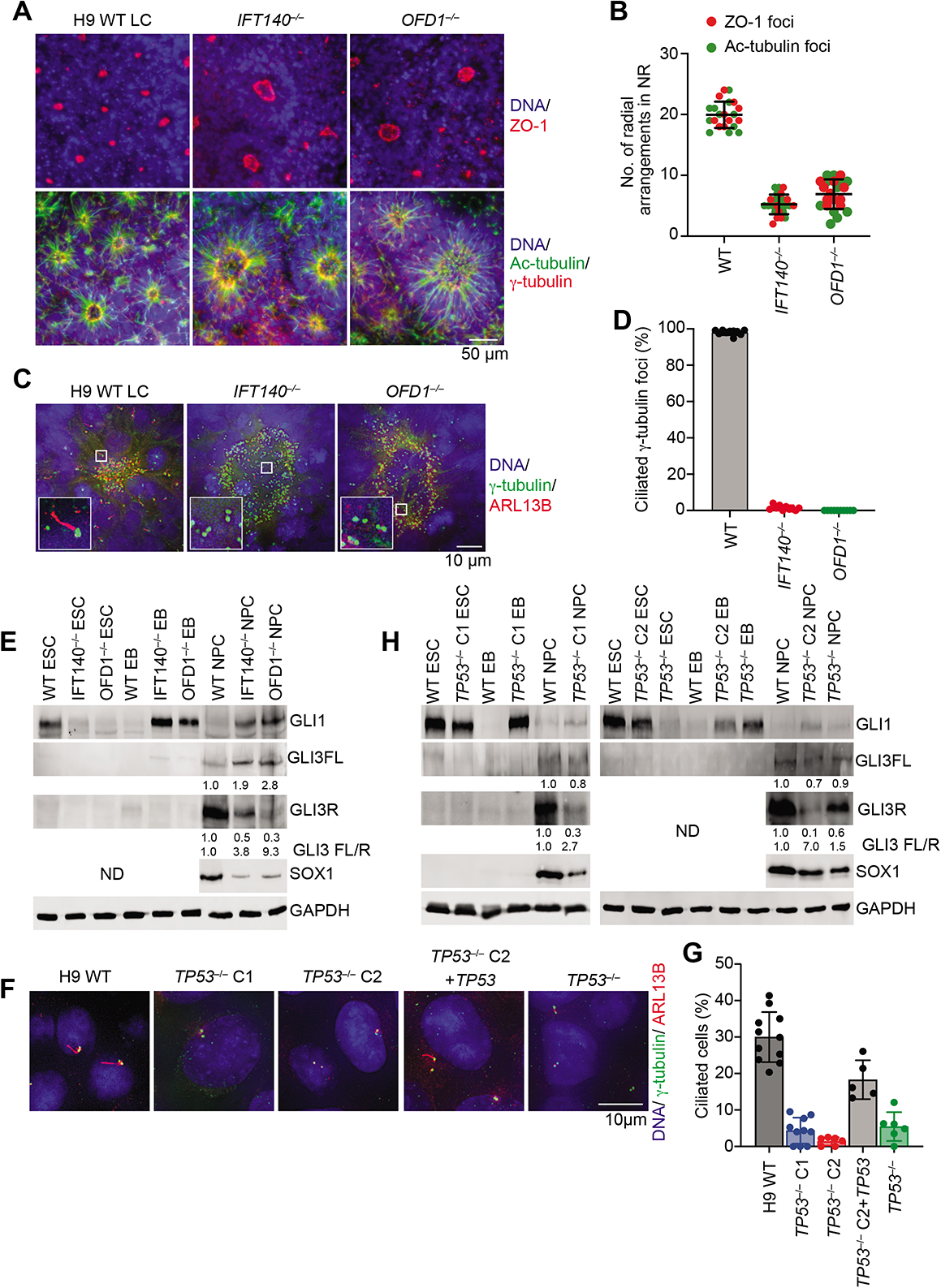
*IFT140^−/−^*, *OFD1^−/−^*, and *TP53^−/−^* hESCs exhibit deficient ciliogenesis, unrestrained Shh signaling, and defective NPC differentiation. (A) Immunofluorescence (IF) images of WT, *IFT140^−/−^*, and *OFD1^−/−^* neural rosettes (NRs) stained with DAPI for DNA and the indicated antibodies. Top and bottom panels show different NRs of each cell line. (B) Quantification of the number of radial arrangements (as defined by ZO-1 and Ac-tubulin foci) within each WT or the indicated mutant NR. Each dot represents one NR. Mean ± s.d.; n=10 for both ZO-1 and Ac-tubulin. (C) Immunofluorescence (IF) images of WT, *IFT140^−/−^*, and *OFD1^−/−^* NRs stained with DAPI for DNA (blue) and antibodies against ARL13B (red) and γ-tubulin (green). (D) Quantification of the percentage of ciliated γ-tubulin foci in each NR. Mean ± s.d.; n=10. (E) Western blots of lysates of H9 WT,H9 *IFT140^−/−^*, and H9 *OFD1^−/−^* ESCs, EBs, and NPCs with the indicated antibodies. (F) (A) Immunofluorescence (IF) images of WT ESCs, *TP53^−/−^* ESCs, and *TP53^−/−^* ESCs re-expressing the *TP53* transgene stained with DAPI for DNA (blue) and antibodies against ARL13B (red) and γ-tubulin (green). (G) Quantification of the percentage of ciliated cells in (F). Each dot represents one field of view. Mean ± s.d.; n=11 (total 695 cells) for WT; n=10 (total 237 cells) for *TP53****^−/−^*** C1; n=6 (total 268 cells) for *TP53****^−/−^*** C2; n=5 (total 183) for *TP53****^−/−^*** C2 re-expressing *TP53*; n=6 (total 290) for *TP53*^−/−^. (H) Western blots of lysates of WT and *TP53^−/−^* ESCs, EBs and NPCs with the indicated antibodies.

The GLI proteins are effectors of Hedgehog pathway and primary cilia are required for formation of both GLI activator and repressor forms (Bangs and Anderson, 2017). Mouse mutants of IFT proteins disrupt primary cilia formation and inhibit GLI3 processing to its repressor form (Huangfu and Anderson, 2005; Liu et al., 2005; May et al., 2005). OFD1 mutant mouse embryonic stem cell lines lack primary cilia and display reduced processing of GLI3 full length to GLI3R during early differentiation (Hunkapiller et al., 2011). Since *IFT140^−/−^/OFD1^−/−^* hESCs and NRs cannot form primary cilia, we examined status of Shh signaling through GLI protein kinetics during differentiation to NPCs. Shh signaling was downregulated during the differentiation of WT hESCs to NPCs, as evidenced by decreased protein levels of GLI1 and full-length GLI3 and accumulation of GLI3R, the cleaved repressor form of GLI3 (Figure 5E). Consistent with well-established roles of primary cilia in regulating Shh signaling, *IFT140^−/−^* and *OFD1^−/−^* NPCs (which lacked primary cilia) had elevated levels of GLI1, 1-2 fold increase in full length GLI3 and 50-70% decrease in levels of GLI3R (Figure 5E), indicative of the failure to downregulate Shh signaling. Increase in GLI1 protein levels in the absence of primary cilia in *IFT140^−/−^/OFD1^−/−^* EBs and NPCs suggest that this may be a cilia-independent effect. Since primary cilia are required for GLI3 processing, reduced processing of GLI3 full length to GLI3R in *IFT140^−/−^/OFD1^−/−^* NPCs is most likely a cilia-dependent effect. Thus, during early hESC differentiation, primary cilia are required to form organized NRs where GLI3 is processed to GLI3R to allow SOX1 expression and NPC differentiation.

### BBS9 is a TP53 target gene that regulates ciliogenesis and NPC differentiation

Our CRISPR-Cas9 screen suggests that genes involved in ciliogenesis and Shh signaling are major regulators of NPC differentiation. We tested whether the NPC differentiation defects of *TP53^−/−^* hESCs were linked to defective ciliogenesis and dysregulated Shh signaling. *TP53^−/−^* hESCs were deficient in ciliogenesis (Figures 5F and 5G). This defect was partially rescued by re-expressing the WT *TP53* transgene. As described above, *TP53^−/−^* hESCs formed disorganized NRs during NPC differentiation. These NRs displayed reduced cilia formation (Figures S9A and S9B). Similar to *IFT140^−/−^/OFD1^−/−^* cells, *TP53^−/−^* EBs and NPC-like cells exhibited increased *GLI1* mRNA and GLI1 protein levels and decreased GLI3R levels (Figures 5H and S9C), consistent with unrestrained Shh signaling. *TP53^−/−^* NRs exhibit 40-50% decrease in primary cilia formation and this may account for partial processing of GLI3 full length to GLI3R (Figure 5H). Thus, ciliogenesis and Shh signaling are indeed perturbed in *TP53^−/−^* hESCs during differentiation.

We then sought to determine whether transcriptional targets of TP53 regulated ciliogenesis and Shh signaling. TP53 occupies neurogenesis genes in the stem cell state, suggesting that it might promote commitment to neural lineages at the onset of differentiation. Among common DEGs between WT and *TP53^−/−^* hESCs, 18 of them contained TP53 peaks in the ChIP-seq data of both hESCs and NPCs (Figure 6A), suggesting that they might be direct TP53 target genes throughout the differentiation process. Indeed, the list of these 18 genes included the best characterized TP53 target gene *CDKN1A* (Figure 6B), which encodes a cyclin-dependent kinase inhibitor that arrests the cell cycle in response to various stimuli (Abbas and Dutta, 2009). Strikingly, the list also included the Bardet-Biedl syndrome 9 (*BBS9*) and patched domain-containing protein 4 (*PTCHD4*) genes. *BBS9* encodes a core subunit of the BBSome, which regulates IFT assembly and transport within the cilia (Wei et al., 2012). *PTCHD4* encodes a negative regulator of Shh signaling with homology to PTCH1 (Chung et al., 2014). *BBS9* and *PTCHD4* are thus candidate TP53 target genes with direct functions in ciliogenesis and Shh signaling.

**Figure 6.**
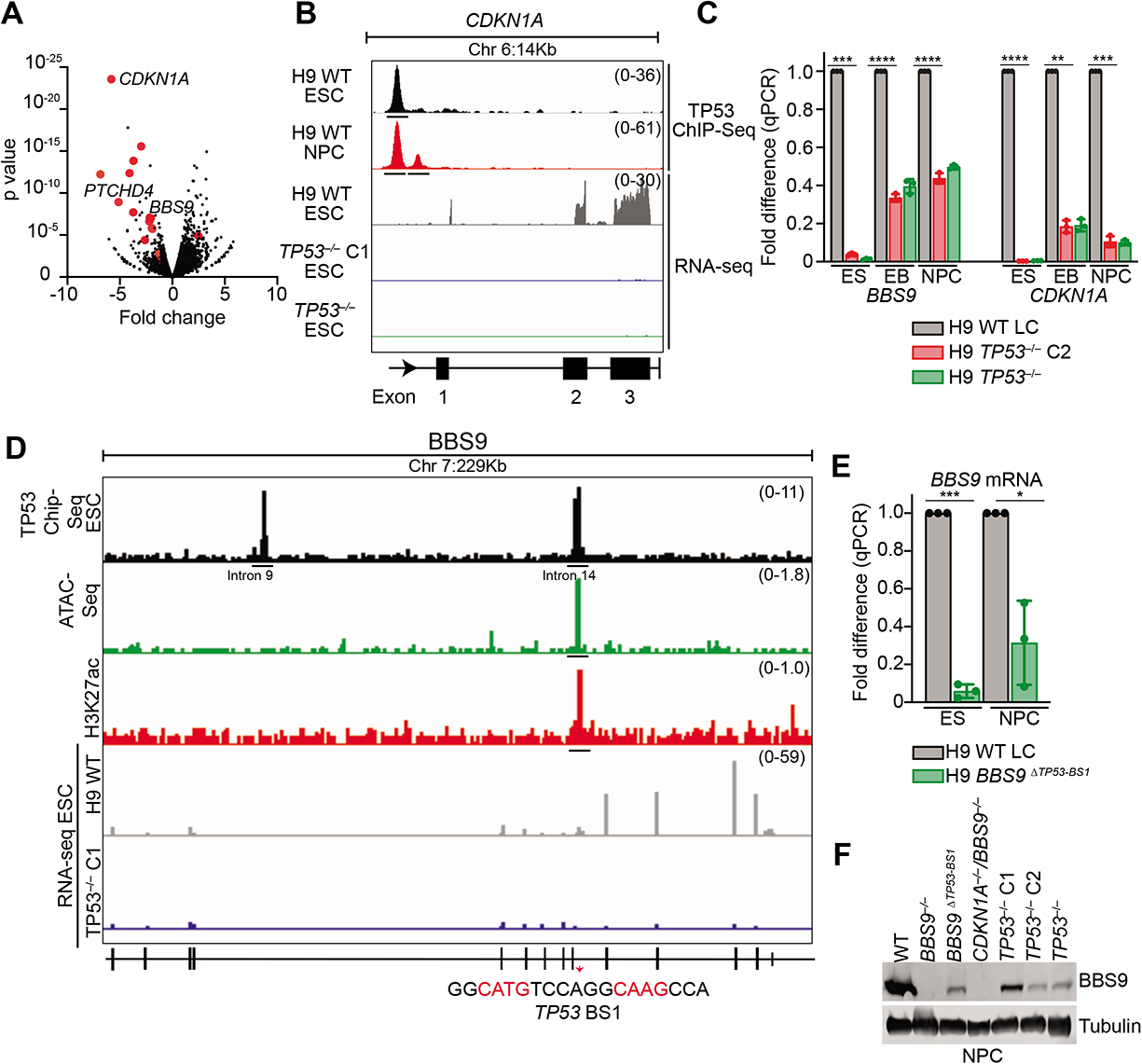
*BBS9* is a TP53 target gene. (A) Volcano plot of RNA-seq data of WT and *TP53****^−/−^*** ESCs showing the log2 fold change and p value of individual genes. DEGs with TP53 ChIP-seq peaks in their vicinity are shown as red dots. (B) TP53 ChIP-seq and RNA-seq tracks from the Integrative Genomics Viewer in the *CDKN1A* gene in WT hESCs and NPCs and in *TP53****^−/−^*** hESCs. The TP53 ChIP-seq peaks in the *CDKN1A* promoter are underlined. (C) qPCR analysis of the mRNA levels of *BBS9* and *CDKN1A* in WT and *TP53****^−/−^*** hESCs, embryoid bodies (EBs), and NPCs. Mean ± s.d.; n = 3 independent experiments; ** p<0.01, *** p<0.001, **** p<0.0001. (D) TP53 ChIP-seq and RNA-seq tracks from the Integrative Genomics Viewer in the *BBS9* gene in WT hESCs and NPCs and in *TP53****^−/−^*** hESCs. The TP53 ChIP-seq peaks in introns 9 and 14 are underlined. Previously published data show that the TP53 ChIP-seq peak in intron 14 overlaps with H3K27ac ChIP-seq peak and that region in intron 14 shows increased chromatin accessibility. (E) qPCR analysis of the mRNA levels of *BBS9* in WT and *BBS9*^ΔTP53-BS1^ hESCs and NPCs. Mean ± s.d.; n = 3 independent experiments; * p<0.01, *** p<0.001. (F) Western blots of lysates of WT, *BBS9^−/−^*, *BBS9***^ΔTP53-BS1^,** *CDKN1A^−/−^/BBS9^−/−^, TP53^−/−^* NPCs with the indicated antibodies.

*PTCHD4* has already been shown to be a TP53 target gene whose overexpression negatively regulates Shh signaling (Chung et al., 2014). We decided to further characterize *BBS9*. Quantitative RT-PCR confirmed that, similar to *CDKN1A*, expression of *BBS9* was indeed decreased in *TP53^−/−^* hESCs, EBs, and NPCs (Figure 6C). Based on ChIP-seq, TP53 binds to introns 9 and 14 within the *BBS9* gene. The TP53 binding site (TP53 BS1) in intron 14 contains the consensus TP53 binding sequence. From published ChIP-seq and ATAC-seq data, we found that TP53 BS1 in intron 14 overlaps with an enhancer mark (H3K27ac) and shows increased chromatin accessibility (Figure 6D) (GSM2386581, GSM1521726) (Ziller et al., 2015). These findings indicate that TP53 promotes the expression of full length *BBS9*.

To understand functions of *BBS9* and its regulation by *TP53* during NPC differentiation *in vitro*, we generated H9 *BBS9^−/−^* hESCs and a mutant BBS9 hESC line deleted of TP53 BS1 (H9 BBS9^ΔTP53-BS1^) (Figure S10A). Because of the intimate connection between cell proliferation and differentiation, we also created hESCs with *CDKN1A* deleted or with both *BBS9* and *CDKN1A* deleted. Deletion of TP53 binding site in BBS9 resulted in decreased mRNA expression and protein levels of full length BBS9 in both ESCs and differentiated NPCs (Figures 6E and 6F). *In vitro* differentiated *TP53^−/−^* NPCs also showed reduced BBS9 full length protein levels (Figure 6F). Both *BBS9^−/−^* and BBS9^ΔTP53-BS1^ hESCs displayed reduced ciliogenesis (Figures 7A and 7B). During differentiation, *BBS9^−/−^* hESCs and BBS9^ΔTP53-BS1^ hESCs formed disorganized NRs with defective ciliogenesis (Figures 7C, 7D, S10B, S10C). In EBs and NPCs, *BBS9^−/−^* and BBS9^ΔTP53-BS1^ exhibited elevated GLI1 protein levels, decreased GLI3R and SOX1 levels (Figure 7E). Thus, TP53 promotes expression of full length *BBS9* to regulate ciliogenesis and sonic hedgehog signaling during differentiation of hESCs into NPCs.

**Figure 7.**
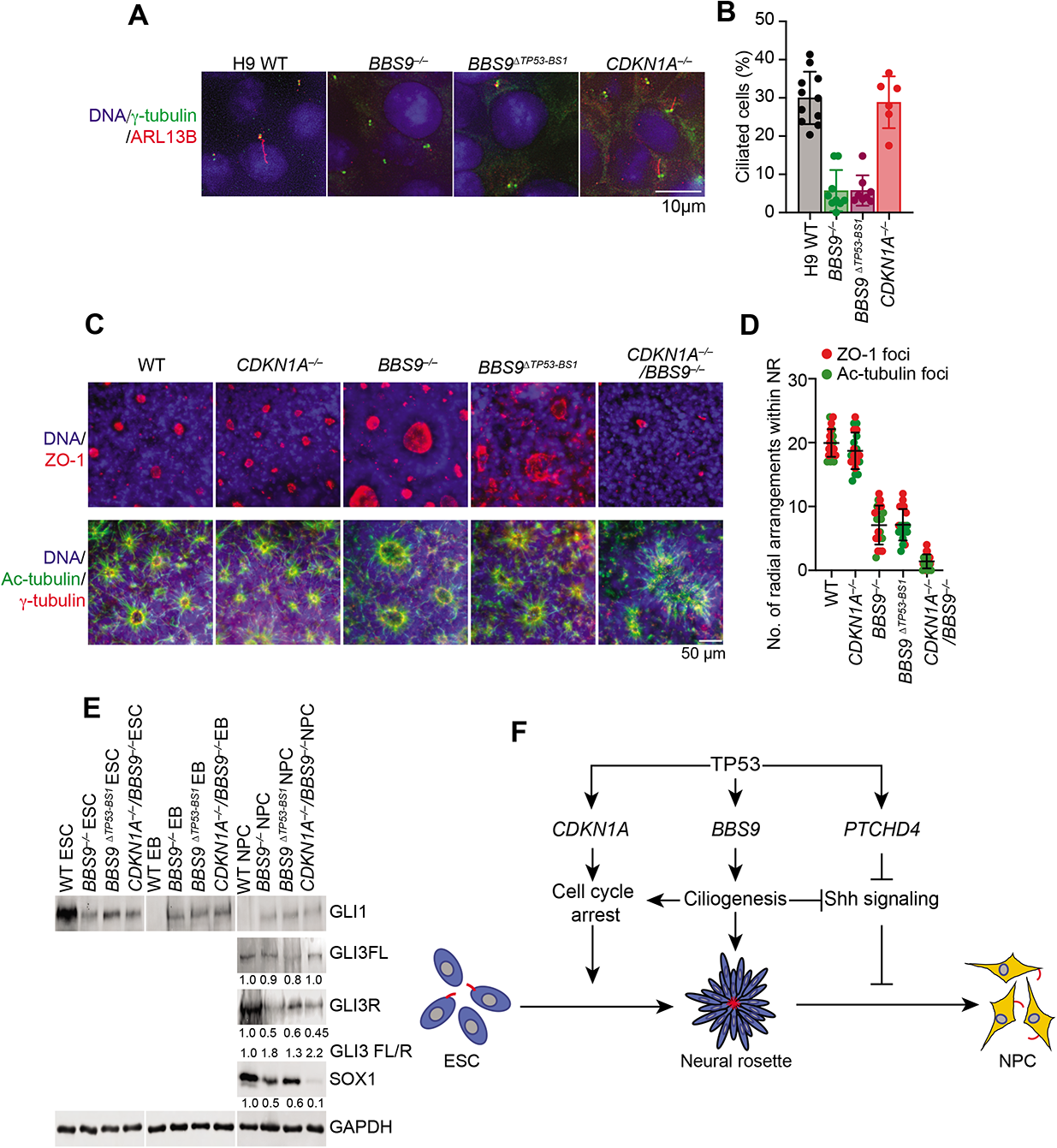
The TP53 target gene *BBS9* promotes ciliogenesis and NPC differentiation. (A) IF images of WT, *BBS9^−/−^*, *BBS9*^ΔTP53-BS1^ and *CDKN1A^−/−^* ESCs stained with DAPI for DNA (blue) and antibodies against ARL13B (red) and γ-tubulin (green). (B) Quantification of the percentage of ciliated cells in (A). Each dot represents one field of view. Mean ± s.d.; n=11 (total 695 cells) for WT; n=9 (total 464 cells) for *BBS9****^−/−^***; n=8 (total 639 cells) for *BBS9*^ΔTP53-BS1^, n=6 (total 542 cells) for *CDKN1A****^−/−^***. (C) Immunofluorescence (IF) images of WT, *CDKN1A^−/−^*, *BBS9****^−/−^***, *BBS9*^ΔTP53-BS1^, and *CDKN1A^−/−^*/*BBS9^−/−^* neural rosettes (NRs) stained with DAPI for DNA and the indicated antibodies. Top and bottom panels show different NRs of each cell line. (D) Quantification of the number of radial arrangements (as defined by ZO-1 and Ac-tubulin foci) within each WT or the indicated mutant NR. Each dot represents one NR. Mean ± s.d; n=10 for ZO-1 and Ac-tubulin. (E) Western blots of lysates of WT, *BBS9****^−/−^***, *BBS9*^ΔTP53-BS1^, and *CDKN1A^−/−^*/*BBS9^−/−^* ESCs, EBs and NPCs with indicated antibodies. (F) Functions of TP53 during hESC differentiation into NPCs. TP53 activates the expression of *CDKN1A* and *BBS9* to execute cell cycle arrest and promote ciliogenesis, respectively. Both processes ensure the formation of tightly compacted neural rosettes, which exert spatiotemporal control of Shh signaling. TP53 also activates the expression of *PTCHD4*, an inhibitor of Shh signaling. Collectively, these mechanisms suppress Shh signaling and promote NPC differentiation.

*CDKN1A^−/−^* hESCs expectedly proliferated faster than WT hESCs (Figure S10D), but did not exhibit ciliogenesis defects (Figures 7A, 7B). Deletion of *CDKN1A* alone did not affect NR formation or reduce SOX1 protein levels in NPCs (Figures 7C, 7D, and S10E), indicating that *CDKN1A* is not required for NPC differentiation. When injected into mice, *CDKN1A^−/−^* hESCs formed largely well-differentiated teratomas suggesting that increased proliferation was not sufficient to cause defect in differentiation (Figure S10F). On the other hand, deletion of both *BBS9* and *CDKN1A* exacerbated the NR formation defects caused by single *BBS9* deletion (Figures 7C and 7D). Furthermore, *BBS9^−/−^*/*CDKN1A^−/−^* NPCs had much lower SOX1 expression than *BBS9^−/−^* NPCs (Figure 7E). These results indicated that *CDKN1A* contributes to NPC differentiation in cells lacking *BBS9*. Interestingly, deletion of both *BBS9* and *CDKN1A* did not cause more severe ciliogenesis defects than *BBS9* single deletion did (Figures 7C and 7D), suggesting that *CDKN1A* promotes NPC differentiation in pathways independent of ciliogenesis.

## DISCUSSION

During the course of studying genomic instability in teratomas, we unexpectedly discovered a requirement for TP53 in the differentiation of hESCs. Genetic analyses link TP53 to ciliogenesis and Shh signaling during hESC differentiation into NPCs and identify *BBS9* as a critical TP53 target gene in this process.

### Functions of TP53 in differentiation

*TP53^−/−^* hESCs exhibit mitotic defects, increased cell proliferation and develop structural aneuploidy in culture. To understand how TP53 regulates early development, we turned to the teratoma assay. Recent work has shown the teratoma assay to be a promising platform to study early human development. The teratoma model recapitulates multi-lineage development and heterogeneity within cells of the teratoma is comparable to similar fetal cell types (McDonald et al., 2020). We found that, despite proper expression of stem cell markers, *TP53^−/−^* hESCs formed highly immature teratomas with high nuclear cytoplasmic (NC) ratio, increased mitotic index, and reduced ciliogenesis. This finding was unexpected, as two previous studies had reported that TP53-deficient hESCs could form mature teratomas (Merkle et al., 2017; Song et al., 2010). The underlying reasons for this discrepancy are unclear, but could be due to multiple factors. First, the extent of TP53 inactivation varies among studies. In one of the two previous studies (Merkle et al., 2017), hESCs harboring spontaneous point mutations of TP53 were examined. These point mutations might not completely inactivate TP53. By contrast, all *TP53^−/−^* hESC lines generated in our study lacked full-length TP53 protein. While some lines expressed truncated TP53 proteins, these fragments lacked the transactivation domain. Thus, the *TP53^−/−^* hESC lines used in our study harbor nearly complete loss-of-function of TP53 mutations. Second, even a nearly complete inactivation of TP53 produces phenotypes of incomplete penetrance and high variance among individual teratomas (Figure 2B). The limited numbers of TP53-deficient cell lines and teratomas used in previous studies might have prevented the detection of these phenotypes. The advent of CRISPR-Cas9 allowed us to construct multiple *TP53^−/−^* cell lines and produce tens of teratomas. Finally, automated image acquisition enabled broad sampling and rigorous quantification of large histological staining sections of the teratomas in our study. Our results convincingly show that *TP53^−/−^* hESCs are deficient in differentiation and form highly immature teratomas. *In vivo* differentiation assays further confirm a role of TP53 in neural differentiation. Indeed, TP53 occupies neurogenesis genes even in the stem cell state, suggesting that it is poised to promote neurogenesis. On the other hand, the differentiation functions of TP53 are not limited to neural lineages, as TP53 loss causes differentiation defects in other lineages in teratomas.

A prior study found that decreased expression of TP53 using RNAi leads to increased neuronal differentiation and reduced proliferation rate in both human neuroepithelial cells and brain organoids (Marin Navarro et al., 2020). In our study we have completely ablated TP53 expression using CRISPR-Cas9 in the stem cell state itself and then differentiated cells into NPCs. Differences in cell lines and differentiation protocols may account for some of the differences between the two studies.

Depletion of TP53 in *Xenopus laevis* and zebrafish causes defects in embryogenesis (Cordenonsi et al., 2003). Although most *Trp53^−/−^* mice develop to term normally, 25% of female *Trp53^−/−^* mice die during embryogenesis due to defects in neural tube closure (Armstrong et al., 1995). This subtle defect in mice could be more easily distinguishable in our targeted *in vitro* neuronal differentiation assay. Thus, the function of TP53 in early development is evolutionarily conserved. In *Trp53^−/−^* mice, the TP53 homologs, TP63 and TP73, compensate for TP53 loss (Molchadsky et al., 2010), explaining their subtle developmental phenotypes. For unknown reasons, TP63 and TP73 cannot effectively compensate for TP53 loss in hESCs, revealing a requirement for TP53 in early human development.

### Mechanisms by which TP53 promotes differentiation

Although *TP53* deletion does not cause whole-chromosome aneuploidy in hESCs, it induces copy number variations (CNVs) at the chromosome arm level (Figures S4A and S4B) (Soto et al., 2017). Moreover, *TP53^−/−^* hESCs likely harbor additional small genetic alterations and mutations. The genomic instability caused by TP53 loss might contribute to the differentiation defects of *TP53^−/−^* hESCs. On the other hand, multiple cell lines with different CNVs and presumably other genetic alterations exhibit similar differentiation defects. Re-expression of the *TP53* transgene in *TP53^−/−^* hESCs, which does not revert the genetic changes, alleviates the differentiation defects. These findings argue against the possibility that differentiation defects of *TP53^−/−^* hESCs are passive, secondary consequences of genomic instability.

Our subsequent findings suggest that TP53 uses multifaceted mechanisms to actively promote hESC differentiation (Figure 7G). TP53 maintains the transcription of *CDKN1A* in all cellular states. As a critical CDK inhibitor, CDKN1A restrains the G1/S transition during the cell cycle to favor differentiation. In addition, TP53 induces the transcription of *BBS9* and *PTCHD4*, whose expression is required for proper ciliogenesis and Shh signaling during NR formation. Collectively, this TP53-orchestrated transcriptional program coordinates cell division, ciliogenesis, and Shh signaling to promote differentiation.

As a conserved microtubule-based organelle, the primary cilium is critical for transducing signals from the external environment to regulate key cellular processes, including cell polarity, differentiation, and proliferation (Goetz and Anderson, 2010). An important function of primary cilia is to regulate Shh signaling. Our genome-wide CRISPR-Cas9 screen reveals a requirement for ciliogenesis genes and negative regulators of Shh signaling in the differentiation of hESCs into NPCs. Inactivation of ciliogenesis genes, including those identified in our screen, causes developmental defects in mouse and other model organisms (Ferrante et al., 2006; Hunkapiller et al., 2011). Their mutations are linked to multi-system developmental syndromes termed ciliopathies in humans (Schmidts et al., 2013; Zaghloul and Katsanis, 2009). Contrary to our screen results, a prior study found that mutant OFD1 mouse stem cell lines exhibit increased Sox1 expression during early differentiation (Hunkapiller et al., 2011). On the other hand, primary cilia and GLI3 processing defects are similar between the OFD1 mutant mouse stem cell line and OFD1^−/−^ hESCs and NPCs in our study. We speculate that differences in SOX1 expression during neuronal differentiation may be due to interspecies variations and needs further study. However, our findings extend the role of ciliogenesis and Shh signaling to the differentiation of hESCs into neural lineages and establish TP53 as a key coordinator of NR formation (Figure 7F).

Only 30% of asynchronous hESCs form primary cilia. Upon induction of neuronal differentiation, primary cilia can be observed on the apical side of NR’s and in majority of NPCs. In the absence of ciliary genes, IFT140/OFD1, primary cilia formation is abolished in hESC, NRs and NPCs. This allowed us to study roles of primary cilia in neural rosette organization, GLI protein behavior and hence Shh signaling during early hESC differentiation into NPCs. Inactivation of *IFT140/OFD1* showed us that primary cilia are required for neural rosette organization. GLI1 protein levels increased by 10-15 fold in *IFT140^−/−^/OFD1^−/−^* EB’s suggesting that this is a cilia-independent effect. Importantly, similar to IFT/OFD1 mutants in mouse, we also found that primary cilia are required for complete processing of GLI3 full length into GLI3R during early human NPC differentiation (Huangfu and Anderson, 2005; Liu et al., 2005; May et al., 2005). Thus, the CRISPR-Cas9 screen provided us with a tool to understand ciliogenesis and Shh signaling during early NPC differentiation. We applied our findings to *TP53^−/−^* cells during NPC differentiation. Inactivation of *TP53* in hESCs causes deficient ciliogenesis in stem cell state itself. Upon induction of NPC differentiation, decreased primary cilia formation in TP53 ablated cells resulted in disorganized NR formation. GLI1 protein levels increase in *TP53^−/−^* EBs and NPCs compared to their respective controls. Since this occurs even in the absence of primary cilia in *IFT140^−/−^/OFD1^−/−^* EB’s and NPC’s we conclude that this is a cilia-independent effect. Importantly, defective cilia formation in *TP53^−/−^* NRs results in reduced GLI3 processing. Attempts to rescue differentiation defect in *TP53^−/−^* cells using Shh inhibitors were not successful. We speculate that this is because of multiple factors. In *TP53^−/−^* cells, decreased BBS9 expression causes aberrant ciliogenesis and this cannot be rescued by Shh inhibitors. Also, we think ciliogenesis and Shh signaling pathways may not be only responsible for the differentiation defect in the absence of TP53. Nevertheless, our current study clearly shows that in the absence of TP53 we observe a failure to restrain Shh signaling resulting in deficient differentiation into NPCs.

The graded hedgehog signaling response controls dorsal-ventral patterning in the developing mouse neural tube and relies on a gradient of GLI activity (Stamataki et al., 2005). Small perturbations of GLI levels can alter the proper development of distinct cell types. NRs with their apical-basal polarity bear resemblance to the nascent neural tube. Ciliogenesis only occurs on the apical side of NRs, forming dense radial arrays of primary cilia with their tips pointing into the lumen. We propose that this high-density, ordered arrangement of cilia ensures proper Shh signaling and cell lineage specification in NRs (Figure 7F). Loss of TP53 or its downstream target genes, such as *BBS9*, reduces the density of primary cilia and perturbs their organization and function. TP53 further inhibits Shh signaling by promoting the expression of *PTCHD4*, which encodes a negative regulator of Shh signaling. TP53 deficiency thus leads to aberrantly elevated Shh signaling and defective NPC formation.

### Tumor suppression by TP53

As a stress-induced transcription factor, TP53 responds to genotoxic and oxidative stresses and activates distinct transcriptional programs to elicit cell cycle arrest, apoptosis, and ferroptosis, thereby suppressing tumorigenesis. Our study identifies a developmentally regulated transcriptional program of TP53 under basal conditions and links TP53 to ciliogenesis and the control of Shh signaling. Perturbation of this program prevents lineage commitment and produces poorly differentiated immature elements. Poor differentiation is a universal feature of malignant tumors of different tissue origins. Ciliogenesis is generally suppressed during tumorigenesis (Goetz and Anderson, 2010). Aberrant activation of Shh signaling has been observed in several human tumors particularly in medulloblastomas and basal cell carcinoma (Briscoe and Therond, 2013). In a subset of medulloblastomas, the lack of primary cilia causes reduced GLI3 repressor formation resulting in abnormal activation of Shh signaling and aggressive tumors (Bangs and Anderson, 2017; Han et al., 2009). These findings raise the intriguing possibility that the differentiation-promoting functions of TP53 might contribute to tumor suppression.

Teratomas grown from hESCs in heterologous hosts recapitulate early events of human embryogenesis and provide a model for understanding human teratomas. Even though *TP53* loss in hESCs produce immature teratomas in the mouse, both mature and immature teratomas in human patients rarely contain *TP53* mutations. Whether the developmentally regulated transcriptional program of TP53 reported herein remains intact in immature human teratomas needs future investigations. *TP53* mutations in human cancers typically do not involve the complete loss-of-function of both alleles. It will be interesting to test whether hESCs engineered to harbor *TP53* missense mutations in human cancers exhibit differentiation defects in teratomas and human organoids. Human Li-Fraumeni syndrome (LFS) patients, who have germ-line *TP53* mutations, are susceptible to developing cancers in multiple tissues (Malkin et al., 1990). Future experiments are needed to test whether differentiation defects of stem/progenitor cells caused by TP53 deficiency contribute to tumorigenesis in LFS patients and, more broadly, whether dysfunction of the developmentally regulated transcriptional program of TP53 contributes to poor differentiation of human cancers driven by TP53 inactivation.

## METHODS

### Cell culture and gene editing

Human embryonic stem cells (hESCs) (WA01-H1 and WA09-H9) were purchased from WiCell Research Institute, Wisconsin, USA. The hESCs were cultured under feeder-free conditions in mTeSR1-cGMP media (STEMCELL Technologies) on plates coated with Matrigel (Corning). Versene (Thermo Fisher Scientific) was used to passage hESCs. The hESCs were routinely tested for mycoplasma to avoid any contamination.

For the generation of *TP53^−/−^* hESCs, sgRNAs targeting exons 2 or 4 of *TP53* were cloned into px458-GFP, px459-puro, or pLenti-Cas9 plasmids (obtained from Addgene)(Ran et al., 2013). The sequences of sgRNAs are: AAGGGACAGAAGATGACAG, TGAAGCTCCCAGAATGCCAG and TCGACGCTAGGATCTGACTG. The px458/px459 plasmids were transfected into hESCs with electroporation using the Lonza 4D-Nucleofector according to the manufacturer’s instructions. One day after electroporation, cells were selected either by flow cytometry (FACS) for GFP expression or by treatment with 1 µg/ml puromycin. Selected cells were allowed to grow into colonies, which were then isolated to obtain the H9 *TP53^−/−^* C1 cell line. A population of selected cells was used for the H1 *TP53^−/−^* cells. For the packaging of lentiviruses, pLenti-Cas9 plasmids (with the *TP53* sgRNAs or without an sgRNA) along with pMD2.G and psPAX2 were transfected into HEK293FT cells using Lipofectamine 2000 (Life Technologies). Viral supernatants were collected at 2-3 days after transfection, concentrated using Lenti X concentrator (Clontech), and added to H9 hESCs growing on Matrigel along with 4 µg/ml polybrene. The infected cells were selected with 1 µg/ml puromycin. The surviving clones were picked to obtain the H9 *TP53^−/−^* C2 cell line. A population of surviving cells was collected and referred to as H9 *TP53^−/−^* to avoid clonal bias. Surviving cells from H9 hESCs infected with the lentivirus without sgRNAs were used as H9 WT lentivirus controls (referred to as H9 WT LC). The loss of TP53 in the knockout cell lines was verified by Western blotting and DNA sequencing of the *TP53* alleles.

H9 *IFT140^−/−^*, *OFD1^−/−^*, *CDKN1A^−/−^*, and *BBS9^−/−^* clonal hESC lines were generated using the lentivirus procedure as described above using the following guides: *IFT140* sgRNA, TACCCCGATCCTTAAACGTC; *OFD1* sgRNA, CCGACATCACCGTGCTCCGT; *CDKN1A* sgRNAs, GGAGAAGATCA-GCCGGCGTT and ACTCTCAGGGTCGAAAACGG; *BBS9* sgRNA, AGTTCTCGAGATTGAACGTT. The *CDKN1A^−/−^* hESC line was further infected with the lentivirus containing sgRNAs targeting *BBS9* (AGTTCTCGAGATTGAACGTT and TGATTCTGCTGATCGCGCTG) to generate the *CDKN1A^−/−^*/*BBS9^−/−^* hESC clonal cell line.

All stem cell work described in this manuscript has been conducted under the oversight of the Stem Cell Research Oversight (SCRO) Committee at University of Texas Southwestern Medical Center (UT Southwestern). When establishing stem cell research standards, UT Southwestern follows the International Society for Stem Cell Research (ISSCR) guidelines for stem cell research and clinical translation and National Institutes of Health guidelines for human stem cell research.

### Cell proliferation assay and flow cytometry

WT, *TP53^−/−^*, and *CDKN1A^−/−^* hESCs were plated in triplicates on Matrigel-coated plates. At 24-h intervals, cells were dissociated, stained with Trypan blue, and counted using a Bio-Rad TC20 automated cell counter. For flow cytometry, hESCs were fixed with 2-4% PFA, permeabilized with methanol, and then stained with anti-SOX2 (APC; Miltenyi Biotec), anti-OCT3/4 (PE; Miltenyi Biotec), and anti-NANOG (BV421; BioLegend) antibodies and DAPI. Cells were analyzed using a Beckman Coulter CytoFLEX flow cytometer.

### Giemsa (G) banding and karyotyping

hESCs and NPCs (differentiated by the 3D embryoid body protocol) were grown in Matrigel-coated or regular 25 cm^2^ tissue culture flasks with the appropriate media. The live cells in culture flasks were shipped to WiCell Research Institute for G banding and karyotyping. For reversine treatment, hESCs were treated with 1 µM reversine for 48 h. After reversine was washed off, the surviving cells were allowed to recover in culture for 2 weeks before being sent to WiCell for G banding analysis.

### Live-cell imaging

hESCs were grown in Matrigel-coated Nunc Lab-Tek chambered coverglass (Thermo Fisher Scientific). Aphidicolin was added at 200 ng/ml for 12 h to synchronize cells in the G1 phase. Before imaging, the cell-permeable Hoechst 33342 dye (Life Technologies) was added to cells at 10-50 ng/ml. Images were captured at 5-min intervals for 10-18 h using a 40X objective and an Evolve 512 Delta EMCCD camera on a Leica inverted microscope equipped with an environmental chamber that controls temperature and CO2. Time-lapsed videos displaying elapsed times between consecutive frames were assembled using the MetaMorph Software (MDS Analytical Technologies) and analyzed using Image J. The mitotic duration between nuclear envelope breakdown and anaphase onset was manually determined for each cell. Mitotic defects were visually identified and counted. GraphPad Prism was used to generate the scatter plots and stacked bar graphs.

### Immunofluorescence

hESCs were grown on Matrigel-coated coverslips. Cells were fixed with the 4% paraformaldehyde solution containing 0.5% Triton X-100. Fixed cells were incubated with the blocking solution (3% BSA in PBS containing 0.1% Tween-20) for 1 h. Cells were then treated with primary antibodies diluted in the blocking solution and incubated overnight at 4°C. Primary antibodies used for staining included anti-ZO-1 (Invitrogen), anti-SOX2 (Abcam), anti-NANOG (Thermo Fisher Scientific), anti-OCT3/4 (Santa Cruz Biotechnology), anti-SOX1 (Cell Signaling), anti-KI67 (Abcam), anti-γ-tubulin (Abcam), anti-acetylated tubulin (Sigma-Aldrich), anti-ARL13B (Proteintech). The slides were then washed with PBS containing 0.1% Tween-20, stained with secondary antibodies and DAPI and mounted with the VECTASHIELD antifade mounting medium (Vector Laboratories). The slides were viewed with a DeltaVision microscope (GE healthcare). Sum stack of image slices was used for intensity quantification. ImageJ was used to determine the signal intensities of nuclei. Normalized signal intensities were calculated by subtracting background signals from nuclear signals.

### Teratoma formation

Human ESCs were injected subcutaneously into 6-8 weeks old female SCID-NOD mice that were obtained from the UT Southwestern Breeding Core. For each injection, 1×10^6^ cells hESCs were resuspended in 1:1 ratio of cold Matrigel:mTeSR1 media. The cell suspension was kept on ice to maintain Matrigel in liquid phase until injection. Each mouse was injected on both flanks subcutaneously with the same hESC line. At least 3 mice were used for each cell line. Mice were sacrificed at 7-9 weeks after the injection and teratomas were isolated.

### Tissue histology and immunohistochemistry

Teratomas were fixed in 10% neutral buffered formalin for 48 h. The fixed tissues were submitted to the Histo Pathology Core at UT Southwestern for processing, embedding, and Hematoxylin and Eosin (H&E) staining. The low and high magnification H&E images of the entire section were captured with a Leica Aperio CS2 slide scanner. Areas of mature and immature elements were manually marked by a pathologist and quantified using ImageJ.

For immunohistochemistry, unstained slides of sectioned tissues were deparaffinized and epitope/antigen retrieval was performed with 10 mM sodium citrate (pH 6.0). The slides were treated with 0.3% H_2_O_2_ to block endogenous peroxidase. The slides were then incubated in 3% BSA and stained with the appropriate primary antibodies, including anti-SALL4 (Abcam), anti-GFAP (Abcam), and anti-Glypican-3 (Abcam). After an overnight incubation in primary antibodies, slides were stained with secondary antibodies and DAPI before mounting with the VECTORSHIELD antifade medium (Vector Laboratories). The slides were viewed with a DeltaVision microscope.

### Quantitative Western blotting

Cell pellets were incubated with 1X SDS sample buffer and then lysed by sonication. Lysates were cleared by centrifugation and analyzed by SDS-PAGE followed by Western blotting. The following primary antibodies were used: anti-p53 DO-1 (Santa Cruz Biotechnology) that recognizes epitopes in the region containing residues 11-25, anti-p53 pAB1801 (Santa Cruz Biotechnology) that recognizes epitopes in the region containing residues 32-79, anti-p53 C-terminal antibody (Sigma-Aldrich) that recognizes epitopes in the region containing residues 334-383, anti-GAPDH (Cell Signaling Technology), anti-OCT3/4 (Santa Cruz Biotechnology), anti-SOX2 (Abcam), anti-NANOG (Santa Cruz Biotechnology), anti-Tubulin (Millipore Sigma) and anti-BBS9 (Millipore Sigma). Secondary antibodies used were anti-rabbit or anti-mouse IgG (H+L) (Dylight 680 or 800 conjugates) (Cell Signaling Technology). The membranes were scanned with the Odyssey infrared imaging system (LI-COR Biosciences).

### Neural differentiation

Human ESCs were differentiated into neural progenitor cells (NPCs) using the STEMdiff neural system (STEMCELL Technologies). Briefly, 2-3 × 10^6^ hESCs suspended in the neural induction medium with the ROCK inhibitor Y27632 were seeded on AggreWell 800 plates (STEMCELL Technologies) for embryoid body (EB) formation. The neural induction medium was changed every day for 5 days. EBs were harvested on day 5 using 37-μm reversible strainers. Collected EBs were replated on fresh Matrigel-coated plates and incubated for 7 days to allow neural rosette (NR) formation. NRs were then selected using STEMdiff NR selection reagent. Selected cells were replated and allowed to grow for another 4-6 days in neural induction medium to form neural progenitor cells (NPCs). NPCs were passaged in STEMdiff neural progenitor medium.

### RNA-seq

RNA was extracted from hESCs and NPCs using TRIzol and isopropanol precipitation. Briefly, the cell pellet was incubated with TRIzol and lysed through pipetting. For RNA extraction from teratoma tissues, the frozen tissues were sonicated in TRIzol. After cell and tissue homogenization, chloroform was added to the samples. The samples were vortexed, incubated for 2 min, and centrifuged at 11,000 g for 15 min at 4°C to remove cellular debris. The supernatant was transferred to a fresh tube, and isopropanol was added to precipitate the RNA. After centrifugation and washes with 70% ethanol, the RNA was resuspended in nuclease-free water. Purified RNA was analyzed to determine the RNA integrity number (RIN). Samples with RINs greater than 8 were used for library preparation.

For library construction, RNAs were isolated from two different teratomas for each WT or *TP53^−/−^* cell line and from biological replicates of hESCs and NPCs cell lines. To make libraries for sequencing, TruSeq Stranded mRNA Library Preparation Kit (Illumina) was used to make the libraries for sequencing according to the manufacturer’s instructions. After verification of the library quality, 75-bp single-end sequencing was performed on 12 multiplexed libraries pooled together in one flow cell using a NextSeq 500 high output sequencer (Illumina).

Raw FASTQ files were analyzed using FastQC (v0.11.2)(Andrews, 2010) and FastQ Screen (v0.4.4) (Wingett and Andrews, 2018), and reads were quality-trimmed using fastq-mcf (ea-utils/1.1.2-806) (Aronesty, 2013). The trimmed reads were mapped to the hg19 assembly of the human genome (University of California at Santa Cruz, version from iGenomes) using STAR v2.5.3a (Dobin et al., 2013). Duplicated reads were marked using Picard tools (v1.127; https://broadinstitute.github.io/picard/). The RNA counts generated from FeatureCounts (Liao et al., 2014) were TMM normalized(Robinson et al., 2010), and differential expression analysis was performed using edgeR (Huang da et al., 2009). Statistical cutoffs of p value ≤ 0.05 and log_2_CPM ≥ 0 and were used to identify statistically significant differentially expressed genes (DEGs). Pathway analysis of genes was performed using the gene set enrichment analysis software (GSEA) (Mootha et al., 2003; Subramanian et al., 2005). DEGs heatmaps were clustered by hierarchical clustering using R (http://www.R-project.org) and generated using the Complex Heatmap package (Gu et al., 2016). Tissue specific enrichment analysis (TSEA, http://genetics.wustl.edu/jdlab/tsea/) was performed on the top 200 DEGs between WT and *TP53^−/−^* teratomas(Wells et al., 2015). Predefined pSI values as provided for the pSI package were used.

### Whole genome sequencing

Genomic DNA from cells or teratomas was isolated using the DNeasy Blood & Tissue kit (Qiagen) as per manufacturer’s instructions. The DNA was submitted to Beijing Genomics Institute (BGI) for library preparation and next-generation sequencing (NGS). For copy number analysis, the sequencing reads were aligned to the human reference genome (hg19, with Y chromosome removed) using BWA (v 0.7.12)(Li, 2013). Duplicate reads were removed from the alignment using Picard tools (c 2.2.1) (http://broadinstitute.github.io/picard/). HMM copy (v 1.18.0) was used to detect copy number variations (Lai D, 2016). A log_2_ ratio of greater than 0.3 was considered a chromosome gain and a log_2_ ratio smaller than −0.3 was considered a chromosome loss.

### Chromatin immunoprecipitation sequencing (ChIP-seq)

hESCs and NPCs (with 10-15×10^6^ cells in each sample) were collected and chemically cross-linked by the addition of 1% formaldehyde for 5 min followed by treatment with 125 mM glycine for 5 min. Cells are then washed twice with PBS and resuspended in the sonication buffer (10 mM Tris HCl pH 7.4, 1 mM EDTA, 0.1% NaDOC, 1% Triton X-100, 0.1% SDS, 0.25% sarkosyl, 1 mM DTT, protease inhibitors from Roche). The cells were sonicated on ice to shear DNA to 300-500 bp. The size of the sheared DNA was confirmed by agarose gel electrophoresis. The whole cell extract was centrifuged for 10 min at 18,000 g at 4°C. The supernatant was transferred to a fresh tube and incubated overnight at 4°C with the p53 antibody (DO-1, Santa Cruz Biotechnology). Samples were then incubated with pre-washed magnetic Dynabeads (Protein G) at 4°C for 2 h. Beads were washed twice with the sonication buffer containing 300 mM NaCl, twice with the sonication buffer containing 500 mM NaCl, twice with the sonication buffer with LiCl (10 mM Tris HCl pH 8.1, 250 mM LiCl, 0.5% NP-40, 0.5% NaDOC, 1 mM EDTA pH 8.0), and twice with the TE buffer (10 mM Tris-HCl, pH 7.5 and 1 mM EDTA pH 8.0). Protein-DNA complexes were eluted from the beads and crosslinking was reversed by overnight incubation in the SDS elution buffer (50 mM Tris-HCl pH 8.1, 1 mM EDTA pH 8.0 and 1% SDS) at 65°C. The immunoprecipitated DNA was treated with RNaseA and proteinase K and recovered using the PCR purification kit (Qiagen). The DNA was then subjected to quality control with Qubit Fluorometer and TapeStation. Libraries from 5-10 ng of ChIP DNA was prepared using the KAPA HTP Library Preparation Kit (Roche) by the Next-Generation Sequencing Core at UT Southwestern. ChIP-seq libraries were sequenced on an Illumina NextSeq 500 using 75 cycle SBS v2 reagents.

Raw FASTQ files were analyzed using FastQC (v0.11.2) (Andrews, 2010) and FastQ_Screen (v0.4.4) (Wingett and Andrews, 2018), and reads were quality-trimmed using fastq-mcf (ea-utils/1.1.2-806) (Aronesty, 2013). The trimmed reads were mapped to the hg19 assembly of the human genome (University of California at Santa Cruz, version from iGenomes) with bowtie2 (version 2.2.3) (Langmead and Salzberg, 2012). The ChIP-seq peaks were called using MACS2 (version 2.0.10) (Zhang et al., 2008), with a q-value threshold of 0.05 and using the random background of ChIP samples as controls.

### CRISPR-Cas9 screen

Human GeCKO v2 library was obtained from Addgene (#1000000048) and amplified according to the provided instructions. The amplified GeCKO libraries A and B were combined at 1:1 ratio to make the complete GeCKO library. HEK 293FT cells were transfected with the GeCKO library plasmids and the viral packaging vectors pMD2.G and psPAX2 using Lipofectamine 2000 (Thermo Fisher Scientific). Lentiviral supernatant was harvested at 48 h and 54 h after transfection and concentrated using the lenti-X concentrator (Clontech) at 4°C for 12-36 h. After incubation, virus-containing media were centrifuged at 1,500g for 45 min at 4°C, and viral particles were resuspended in PBS and stored at −80°C until use.

For viral titer determination, H9 WT ESCs were infected with different amounts of lentivirus. At 48 h after transduction, cells were dissociated using Accutase and re-seeded on Matrigel-coated plates with or without 0.5 µg/ml puromycin. After another 48 h, when non-transduced cells were all killed by puromycin, cells were dissociated from wells with or without puromycin and counted to determine the transduction efficiency. The amounts of viruses that generated 30-50% transduction efficiency (MOI=0.3-1) were used for subsequent CRISPR-Cas9 screen.

Two biological repeats of H9 ESCs were infected with GeCKO v2 lentiviruses at MOI≤1. For 7 days, cells were maintained in mTeSR1-cGMP with puromycin, and a library coverage greater than 500× was ensured during passaging. At day 7 after transduction, puromycin was withdrawn and cells were cultured for one additional day in mTeSR1-cGMP. At day 8, when cells reached ∼80% confluency, cells were collected and resuspended in STEMdiff™ SMADi Neural Induction medium supplemented with the ROCK inhibitor Y27632 and seeded onto Matrigel-coated 6-well plates. Cells were fed daily with STEMdiff™ SMADi neural induction medium and cultured for 7 days. At day 7 of neural induction, the cells were dissociated with Accutase, pooled, and washed with PBS. 6×10^8^ cells were fixed with 30 ml BD Phosflow™ Fix Buffer I (BD Biosciences) for 20 min at room temperature. After being washed twice with PBS, cells were resuspended in 25 ml BD Phosflow™ Perm Buffer III (BD Biosciences) and stored at −20°C.

About 3×10^8^ cells stored in BD Phosflow™ Perm Buffer III were washed twice with 1% PBSA buffer (PBS containing 1% BSA) and then incubated with rabbit anti-SOX1 antibody (Cell Signaling Technology, #4194) at 1:150 dilution for 1 h at room temperature. Cells were washed twice with the PBSA buffer and incubated with the Alexa Fluor® 488 Donkey Anti-rabbit IgG (H+L) Antibody (Thermo Fisher Scientific) at 1:500 dilution for 30 min at room temperature. After being washed twice with the PBSA buffer, cells were resuspended in 8 ml PBSA buffer. Cells were filtered using the 40-µm cell strainer and sorted using BD FACSAria II SORP or BD FACSAria Fusion sorters. The 1% of cell population with the lowest SOX1 signal was collected. Unsorted cells were also similarly treated and collected.

Genomic DNA was extracted from fixed cells as described(Golden et al., 2017) with minor modifications. The sorted cells were resuspended in 600 µl STE lysis buffer (11 mM EDTA pH 8.0, 10 mM Tris-HCl pH 8.0, 100 mM NaCl, 250 µg/ml proteinase K, 0.25% SDS). 6×10^6^ unsorted cells were resuspended in 1 ml STE lysis buffer. Cell lysis was performed by an overnight incubation at 55°C with occasional vortexing. RNase A (1 mg) was then added to each tube and the samples were incubated at 37°C for 1 h while shaking. gDNA extraction was performed with an equal volume of buffer-saturated phenol (pH 7.9), followed by phenol:chloroform:isoamyl alcohol (25:24:1), followed by chloroform. The aqueous phase after chloroform extraction was separated into 300 µl aliquots in 1.5-ml microtubes. 7.5 µg of Glycoblue (Thermo Fisher Scientific) and 900 µl ethanol were added to each tube and the samples were incubated overnight at −80°C. gDNA was precipitated by centrifugation at 18,000 g for 15 min at 4°C, washed with 1 ml 75% ethanol, air dried for 10 min, and resuspended and incubated in 50 ul buffer AE (10 mM Tris-Cl, 0.5 mM EDTA, pH 9.0) at 37°C for about 10 h. The DNA concentration was measured with the Qubit dsDNA BR assay kit (Thermo Fisher Scientific).

Two rounds of PCR were performed for each gDNA sample to amplify gRNA cassettes with Illumina sequencing adapters and indexes (Sanjana et al., 2014; Shalem et al., 2014). In the first round of PCR (PCR-I), 20 cycles of amplification were performed using Herculase II DNA polymerase (Agilent). For sorted cells, all gDNA from a given sample was distributed into three 50 µl reactions. For unsorted cells, 2 µg of gDNA was used in each 50 µl PCR reaction and a total of 90 reactions were performed to maintain 300× coverage of the library. After PCR-I, all PCR product from a given sample was pooled and mix thoroughly. In the second round of PCR (PCR-II), 2.5 µl of PCR-I product was used as template in each 50 µl reaction. 30 reactions and 6 reactions were performed for unsorted and sorted cells, respectively, to maintain library coverage. The PCR cycle number was optimized (ranging from 10-14 cycles) for each sample to produce similar DNA yield. PCR-II product from a given sample was pooled and DNA was purified with AMPure XP beads (Beckman Coulter) according to the manufacturer’s instructions. After being eluted with 40 µl nuclease-free water, all DNA libraries were checked with Agilent BioAnalyzer High Sensitivity DNA Analysis Kit (Agilent). DNA concentration was measured both by Qubit dsDNA HS assay kit (Thermo Fisher Scientific) and by qPCR using a KAPA Library Quantification Kit (Roche) for Illumina platforms. All samples were pooled and sequenced on Illumina NextSeq 500 sequencer with read configuration as 75 bp, single end and about 20-30 million reads per sample.

The FASTQ files were subjected to quality check using FastQC (v0.11.2) and FastQ_Screen (v0.4.4) (Wingett and Andrews, 2018), and adapters were trimmed using an in-house script. The reference sgRNA library sequences for human GeCKO v2.0 (A and B) were downloaded from Addgene (https://www.addgene.org/pooled-library/). The trimmed FASTQ files were mapped to the reference sgRNA library with mismatch option as 0 using MAGeCK (Li et al., 2014). Read counts for each sgRNA were generated and median normalization was performed to adjust for library sizes. Positively and negatively selected sgRNA and genes were identified with MAGeCK using the default parameters.

## Data availability

All raw sequencing files and peaks files generated in this study have been deposited to the NCBI GEO (https://www.ncbi.nlm.nih.gov/geo) under the accession number GSE168587.

## ACKNOWLEDGMENTS

We thank Eunhee Choi and Sean Goetsch for technical assistance with the teratoma assay, and Sergii Kyrychenko and Jay Schneider for assistance with CRISPR-Cas9 gene targeting in hESCs. We also thank members of the McDermott Sequencing Core at UT Southwestern for providing technical assistance with NGS experiments. This study was supported by grants from the Howard Hughes Medical Institute, the National Institutes of Health (1R01GM124096), Cancer Prevention and Research Institute of Texas (RP160667-P2), and the Welch Foundation (I-1441).

## AUTHOR CONTRIBUTIONS

S.S. conceived the study, designed and performed all the experiments except those noted below, analyzed data and wrote the paper. S.Q. performed the genome-wide CRISPR-Cas9 screen and constructed the *IFT140^−/−^* and *OFD1^−/−^* hESC lines. N.C. performed TP53 ChIP-Seq experiments. A.A.S. and C.X. analyzed RNA-seq and CRISPR-Cas9 data. M.K. and C.X. analyzed ChIP-Seq data. A.K. and C.X. analyzed CNV data. B.M.E. assisted with histopathological analysis of teratomas. H.Y. conceived and supervised the study and edited the paper.

## SUPPLEMENTAL FIGURE LEGENDS

**Figure S1. *TP53^−/−^* hESCs express pluripotency markers and exhibit larger cell and nuclear sizes**

(A) CRISPR-Cas9 strategies for inactivating *TP53*. The positions of gRNAs are indicated. Mutations of *TP53* alleles as revealed by DNA sequencing are shown.

(B,C) Immunofluorescence (IF) images of WT and *TP53^−/−^* hESCs stained with DAPI for DNA and the indicated antibodies.

(D,E) Quantification of the nuclear area (D) and the ratio of nuclear to cell areas (E) of WT and *TP53^−/−^* hESCs. Each dot represents one cell. Mean ± s.d.; n=50-100 cells for (D) and n=30-50 cells for (E).

**Figure S2. *TP53^−/−^* hESCs form solid teratomas with fewer cysts**

(A) Images of intact WT and *TP53^−/−^* teratomas grown in immunodeficient mice.

(B) Hematoxylin and eosin (H&E) staining of WT and *TP53^−/−^* teratoma sections.

**Figure S3. *TP53^−/−^* hESCs form immature teratomas with deficient ciliogenesis and high mitotic index**

(A,B) Hematoxylin and eosin (H&E) staining of WT and *TP53^−/−^* teratoma sections at high magnification. Arrowheads indicate cilia. Black dots indicate stunted cilia. Stars indicate mitotic cells.

(C) Immunohistochemistry (IHC) staining of WT and *TP53^−/−^* teratoma sections with DAPI for DNA and the indicated antibodies.

**Figure S4. *TP53^−/−^* teratomas lack whole-chromosome aneuploidy but exhibit altered transcription of brain-specific genes**

(A,B) Genome-wide plot of chromosome copy number variation (CNV) of H9 WT and *TP53^−/−^* hESCs (A) and teratomas (B).

(C) Tissue-specific expression analysis (TSEA) of the top 200 differentially expressed genes (DEGs) between WT and *TP53^−/−^* teratomas. The size of the hexagon is proportional to the number of transcripts in a particular tissue. The heatmap color scheme depicts the p value of Fisher’s exact test.

**Figure S5. *TP53^−/−^* hESCs are deficient in differentiation into neural progenitor cells (NPCs)**

(A) Flow chart of the three-dimensional (3D) embryoid body (EB) neuronal differentiation protocol with representative images of each stage.

(B) Images of WT and *TP53^−/−^* EBs.

(C) Quantification of the areas of WT and *TP53^−/−^* EBs. Each dot represents one EB. Mean ± s.d.; n=60-100.

(D) Images of WT and *TP53^−/−^* NRs.

(E) Quantification of the areas of WT and *TP53^−/−^* NRs. Each dot represents one NR. Mean ± s.d.; n=100.

(F) Immunofluorescence (IF) images of WT and *TP53^−/−^* NRs stained with DAPI for DNA and the anti-PAX6 antibody.

(G) Immunofluorescence (IF) images of WT and *TP53^−/−^* NPCs and *TP53^−/−^* NPCs re-expressing *TP53* that had been stained with DAPI for DNA and the anti-SOX2 antibody.

(H) Quantification of SOX2 staining intensity of cells in (F). Each dot represents one cell. Mean ± s.d.; n=100.

(I) Immunofluorescence (IF) images of WT and *TP53^−/−^* NPCs stained with DAPI for DNA and the anti-NESTIN antibody.

(J) Western blots of lysates of WT and *TP53^−/−^* hESCs, EBs, and NPCs with the indicated antibodies.

**Figure S6. *TP53^−/−^* NPC-like cells proliferate normally and do not exhibit aneuploidy**

(A) Immunofluorescence (IF) images of WT and *TP53^−/−^* NPCs stained with DAPI for DNA and the anti-KI67 antibody.

(B) Quantification of KI67 staining intensity of cells in (A). Each dot represents one cell. Mean ± s.d.; n=75-100.

(C) IF images of astrocytes differentiated from WT and *TP53^−/−^* NPCs stained with DAPI for DNA and antibodies against GFAP and NESTIN.

(D) IF images of neurons differentiated from WT and *TP53^−/−^* NPCs stained with DAPI for DNA and antibodies against MAP2 and ß-III-Tubulin.

(E) Karyotypes of H9 WT and *TP53^−/−^* C2 NPCs by Giemsa (G) banding. Red arrow indicates chromosome 18q deletion.

**Figure S7. Gene expression changes induced by TP53 loss in the NPC and ESC states**

(A) Venn diagram showing the number and overlap of differentially expressed genes (DEGs) in three pairwise comparisons of WT and *TP53^−/−^* NPCs. DEGs were defined as genes with Log counts per million (CPM)>0, absolute fold change (FC) >1 or <–1, and p value <0.05.

(B) Heatmap of normalized counts of the 589 common DEGs between WT and *TP53^−/−^* NPCs. Genes in the neuron differentiation and cell proliferation pathways are indicated.

(C) TP53 ChIP-seq peaks in the ESC and NPC states.

(D) Venn diagram showing the number and overlap of DEGs between WT and *TP53^−/−^* hESCs. DEGs were defined as genes with Log counts per million (CPM)>0, absolute fold change (FC) >1 or <–1, and p value <0.05.

(E) Heatmap of normalized counts of the 106 common DEGs between WT and *TP53^−/−^* hESCs. Genes in the TP53 and cell proliferation pathways are indicated.

(F) GSEA of the 106 common DEGs between WT and *TP53^−/−^* hESCs.

**Figure S8. Deletion of ciliary genes *IFT140* and *OFD1* in hESCs disrupts ciliogenesis**

(A) Volcano plot of RNA-seq data of WT hESCs and NPCs showing the log_2_ fold change and p value of individual genes. The *SOX1* gene is shown as a red dot.

(B) CRISPR-Cas9 strategies for inactivating *IFT140* and *OFD1*. The positions of gRNAs and mutations of *IFT140* and OFD1 alleles are indicated.

(C) Immunofluorescence (IF) images of WT, *IFT140^−/−^*, and *OFD1^−/−^* hESCs stained with DAPI for DNA and antibodies against ARL13B and γ-tubulin.

(D) Quantification of the percentage of ciliated cells in (C). Each dot represents one field of view. Mean ± s.d.; n=11 (total 695 cells) for WT; n=10 (total 510 cells) for *IFT140^−/−^*; n=10 (total 600 cells) for *OFD1^−/−^*.

(E) Hematoxylin and eosin (H&E) staining of WT, *IFT140^−/−^* and *OFD1^−/−^* teratoma sections at low magnification. Four teratomas in each group were analyzed with representative images shown.

(F) IHC staining of WT, *IFT140^−/−^*, and *OFD1^−/−^* teratoma sections with DAPI for DNA and antibodies against ARL13B and γ-tubulin.

**Figure S9. TP53 deletion affects ciliogenesis within neural rosette and increases GLI1 mRNA levels.**

(A) Immunofluorescence (IF) images of WT and *TP53^−/−^* NRs stained with DAPI for DNA (blue) and antibodies against ARL13B (red) and γ-tubulin (green).

(B) Quantification of the percentage of ciliated γ-tubulin foci in each NR. Mean ± s.d.; n=10.

(C) qPCR analysis of *GLI1* mRNA levels in WT and *TP53****^−/−^*** ESCs and embryoid bodies (EB) during differentiation. Mean ± s.d.; n=4; ** p<0.01.

**Figure S10. The TP53 target gene *BBS9* promotes ciliogenesis**

(A) CRISPR-Cas9 strategies for deleting *BBS9*, TP53-BS1 region within *BBS9*, *CDKN1A*, and *CDKN1A/BBS9*. The positions of gRNAs and mutations of *BBS9* and *CDKN1A* alleles are indicated.

(B) IF images of WT, *BBS9^−/−^*, and *CDKN1A^−/−^*/*BBS9^−/−^* neural rosettes (NRs) stained with DAPI for DNA (blue) and antibodies against ARL13B (red) and γ-tubulin (green).

(C) Quantification of the percentage of ciliated γ-tubulin foci in each NR in (C). Mean ± s.d.; n=10.

(D) Cell proliferation assays of WT and *CDKN1A^−/−^* ESCs. Mean ± s.d.; n=3 independent experiments.

(E) Western blots of lysates of WT and *CDKN1A^−/−^* NPCs with the indicated antibodies.

(F) Hematoxylin and eosin (H&E) staining of WT and *CDKN1A^−/−^* teratoma sections at low magnification. Three teratomas in each group were analyzed with representative images shown.

---

**Table S1.** Karyotypes of *TP53*^−/−^ hESCs and NPCs

| Cell lines | Clonal abnormalities | Nonclonal abnormalities |
| --- | --- | --- |
| H9 WT ESC | 46,XX | — |
| H9 *TP53*<sup>-/-</sup> C1 ESCs | 46,XX, i(20)q(10)[19] | 46,XX, dup (1)(q32q42),<br>i(20)q(10) |
| H9 *TP53*<sup>-/-</sup> C1 ESCs<br>(passage P21) | 46,XX, i(20)(q10)[16]/<br>46,XX, i(20)(q10), dup(21)(q21q22)[3] | 46,XX, t(10;20)(q24;p12) |
| H9 *TP53*<sup>-/-</sup> C2 ESCs | 46, XX, del(18)(q21.3)[13]/ 46,XX[6]<br>28% polyploidy | 47,XX, +12, del (18)(q21.3) |
| H9 *TP53*<sup>-/-</sup> C2 ESCs<br>(passage P21) | 46,XX, del(18)(q21.3)[15]/ 46,XX[3] | 46,XX, t(2;12)(p13;q22),<br>del(18)(q21.3)<br>46,XX,t(3;12)(p11;q24.2),<br>del(18)(q21.3) |
| H9 *TP53*<sup>-/-</sup> ESCs | 47,XX,+X[2]/46XX,i(20)(q10)[2]/ 46,<br>XX [16] | — |
| H9 *TP53*<sup>-/-</sup> ESCs<br>treated with reversine | 46,XX, i(20)(q10)[14]/<br>46,XX[6] | — |
| H9 LC NPC | 46,XX | — |
| H9 *TP53*<sup>-/-</sup> C2 NPCs | 46~47,XX, del (18)(q21.3)[cp2]/<br>46,XX [16] | 47,XX,+9;<br>46,XX,add(2)(p25) |
| H1 *TP53*<sup>-/-</sup> ESCs | 46,XY,t(5;18) (p15.1;q21.1)[4]/<br>46,XY[16] | — |
| H1 *TP53*<sup>-/-</sup> NPCs | 46,XY, t(5;18)(p15.1;q21.1)[3]/<br>46, XY[15] | 46,XY,del (1)(p32);<br>46,XY,add (18)(q23) |

