## Supplementary figures and images for "TP53 promotes lineage commitment of human embryonic stem cells through ciliogenesis and sonic hedgehog signaling"

### Supplement Figures

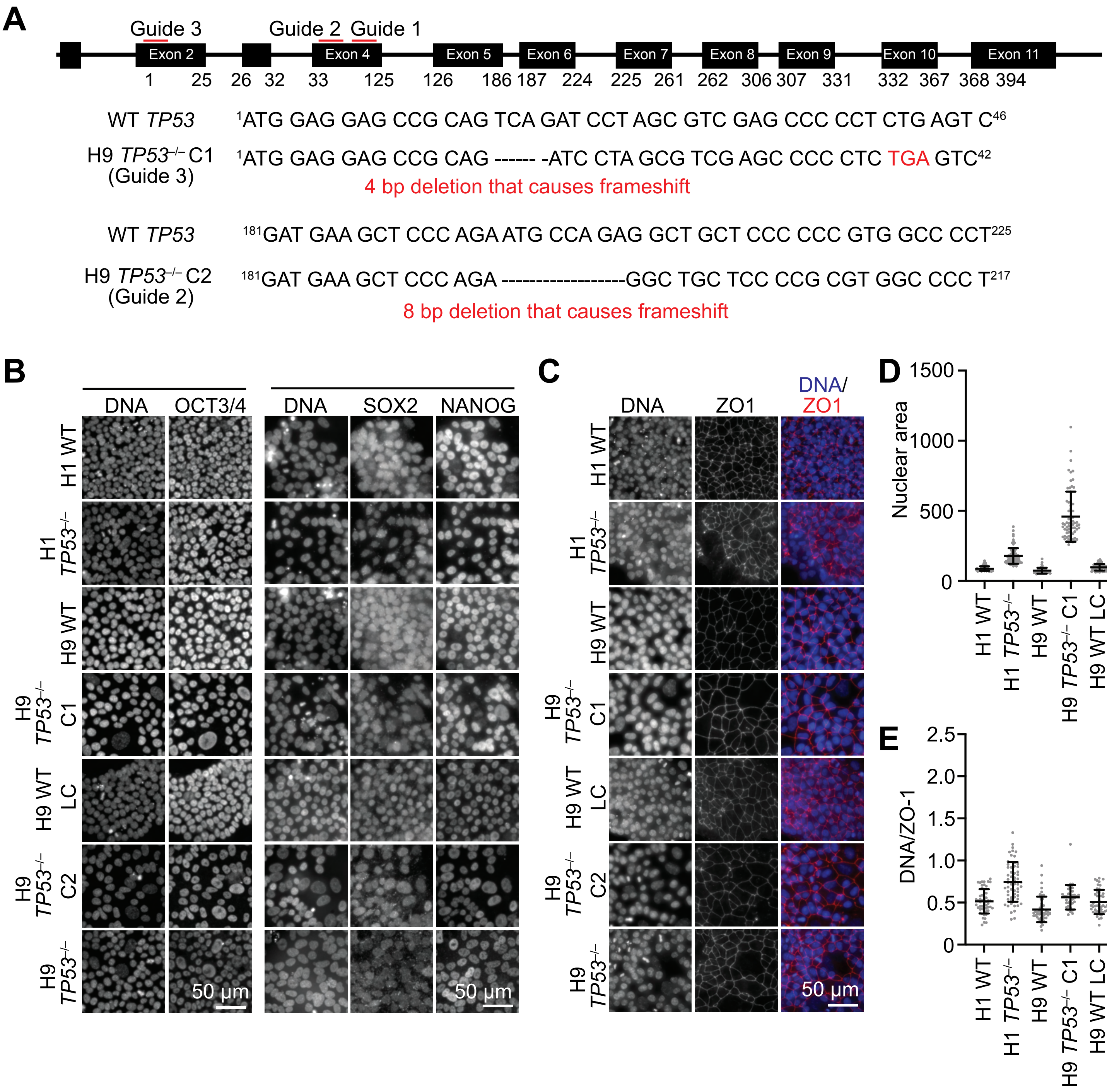

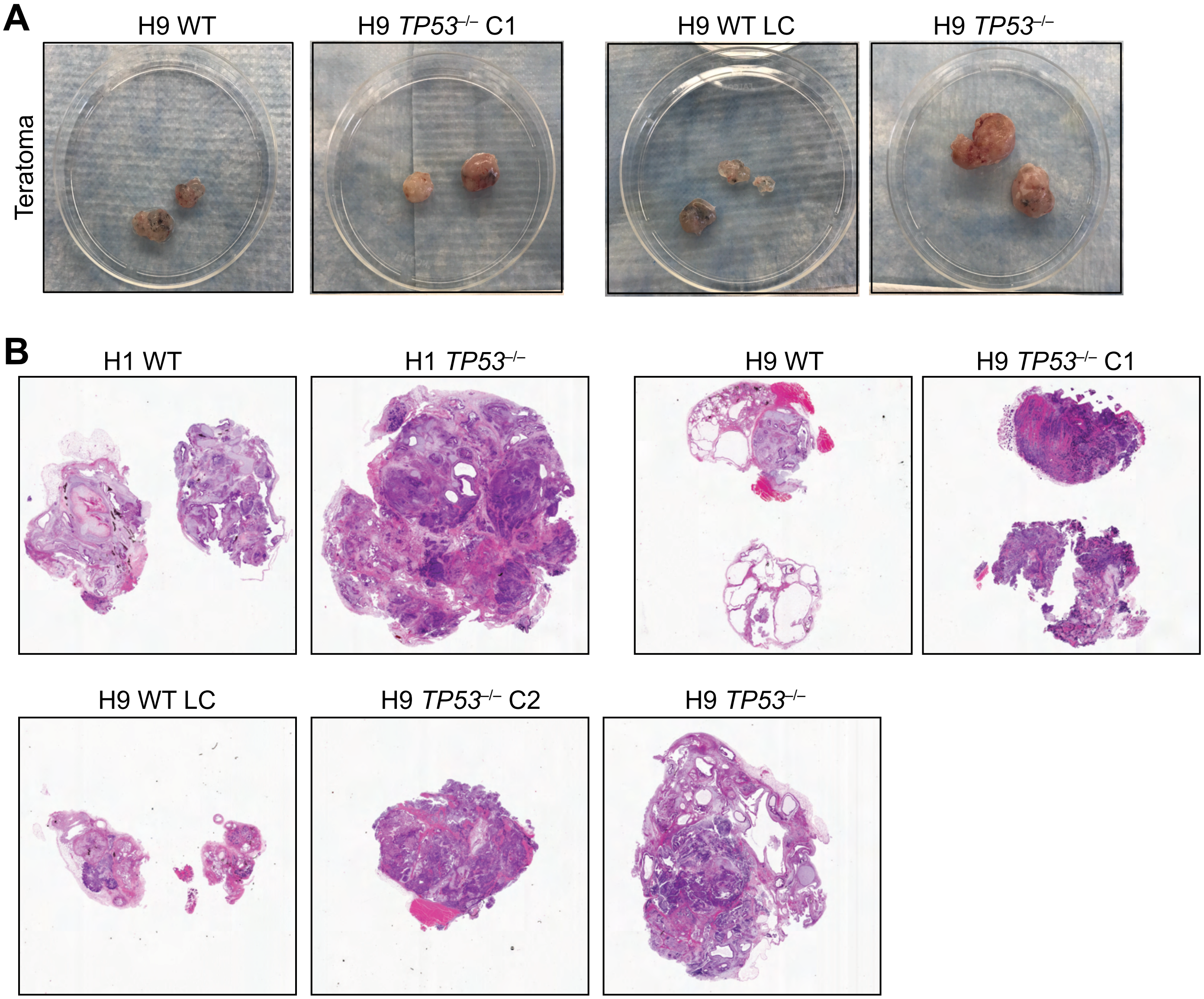

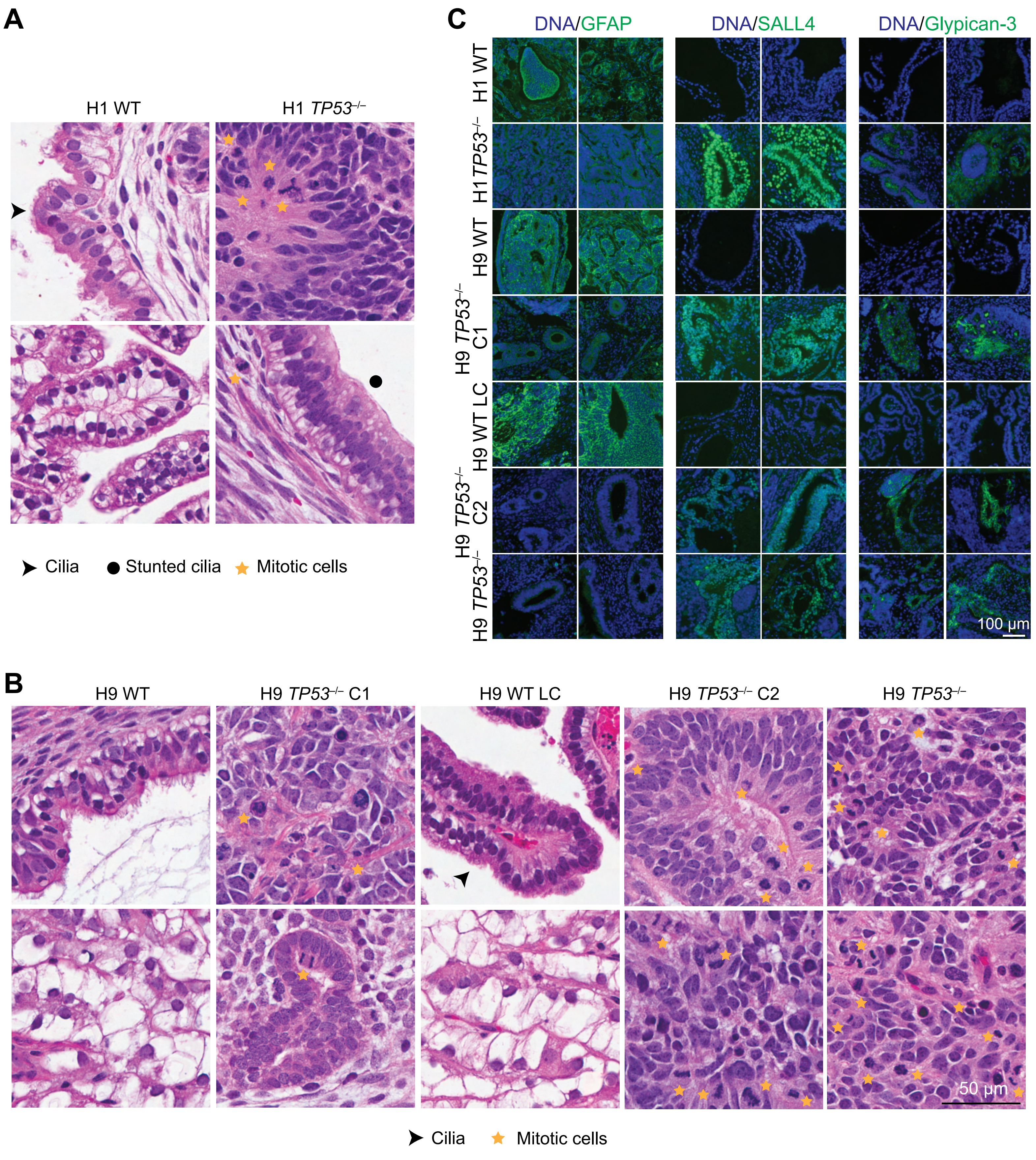

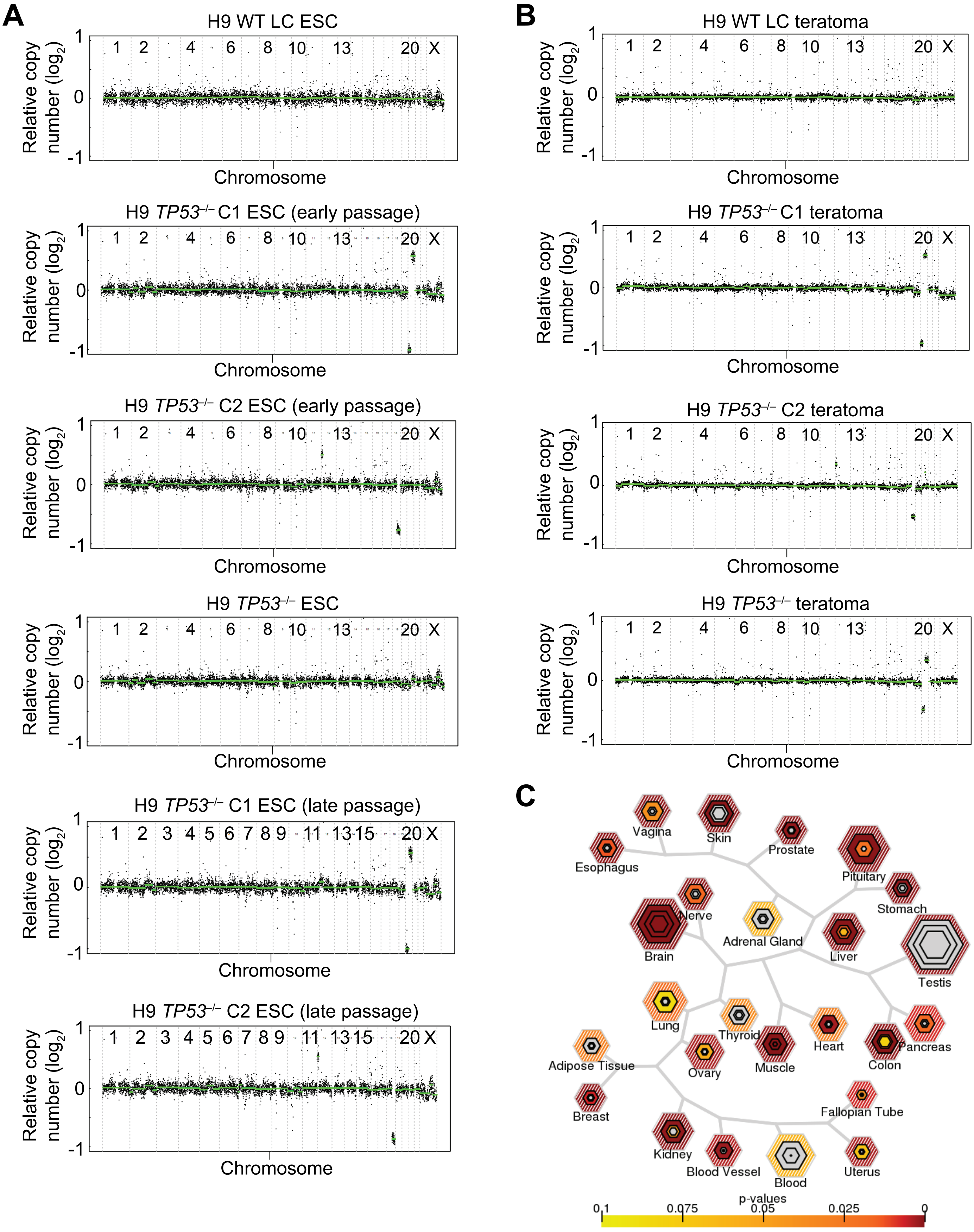

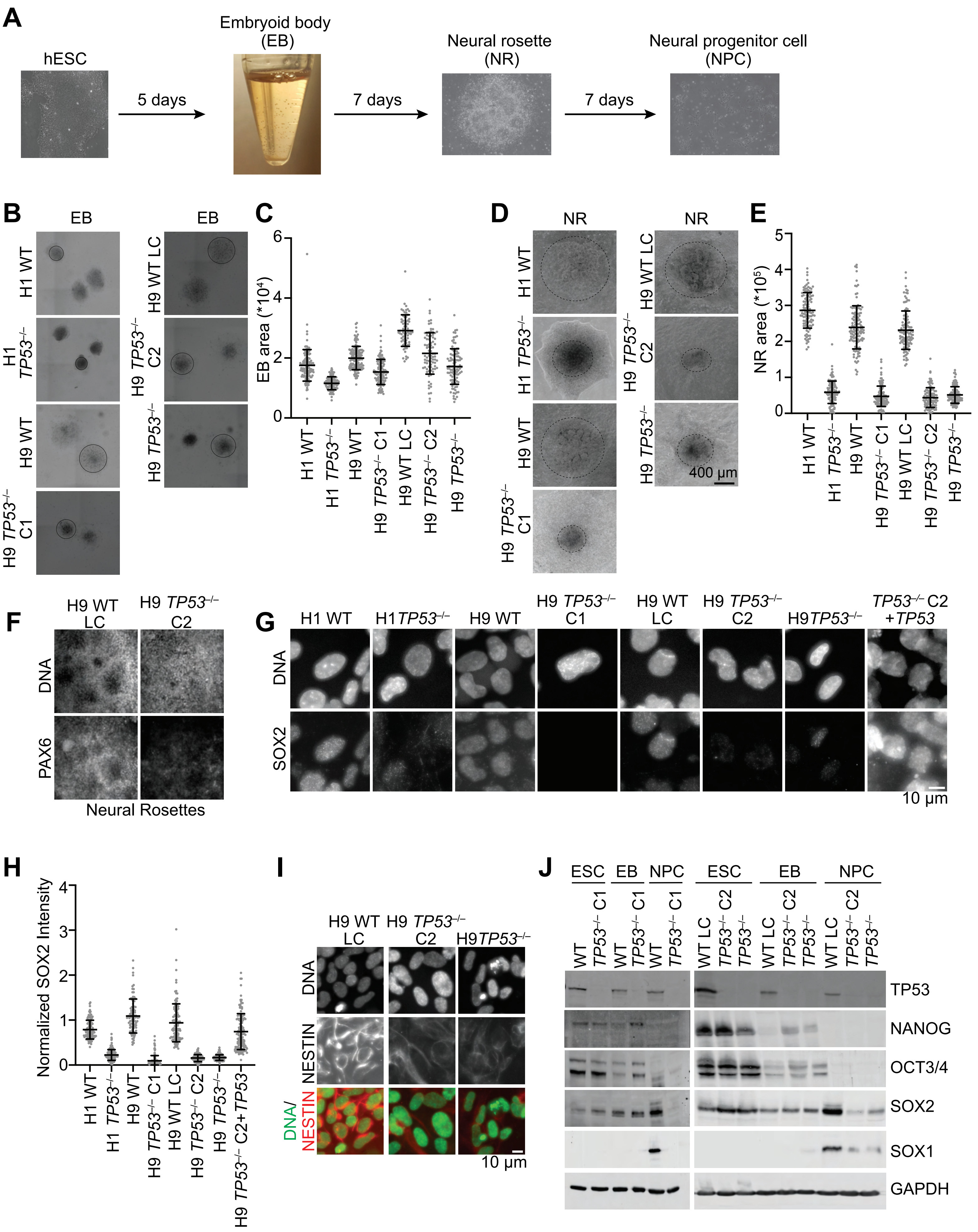

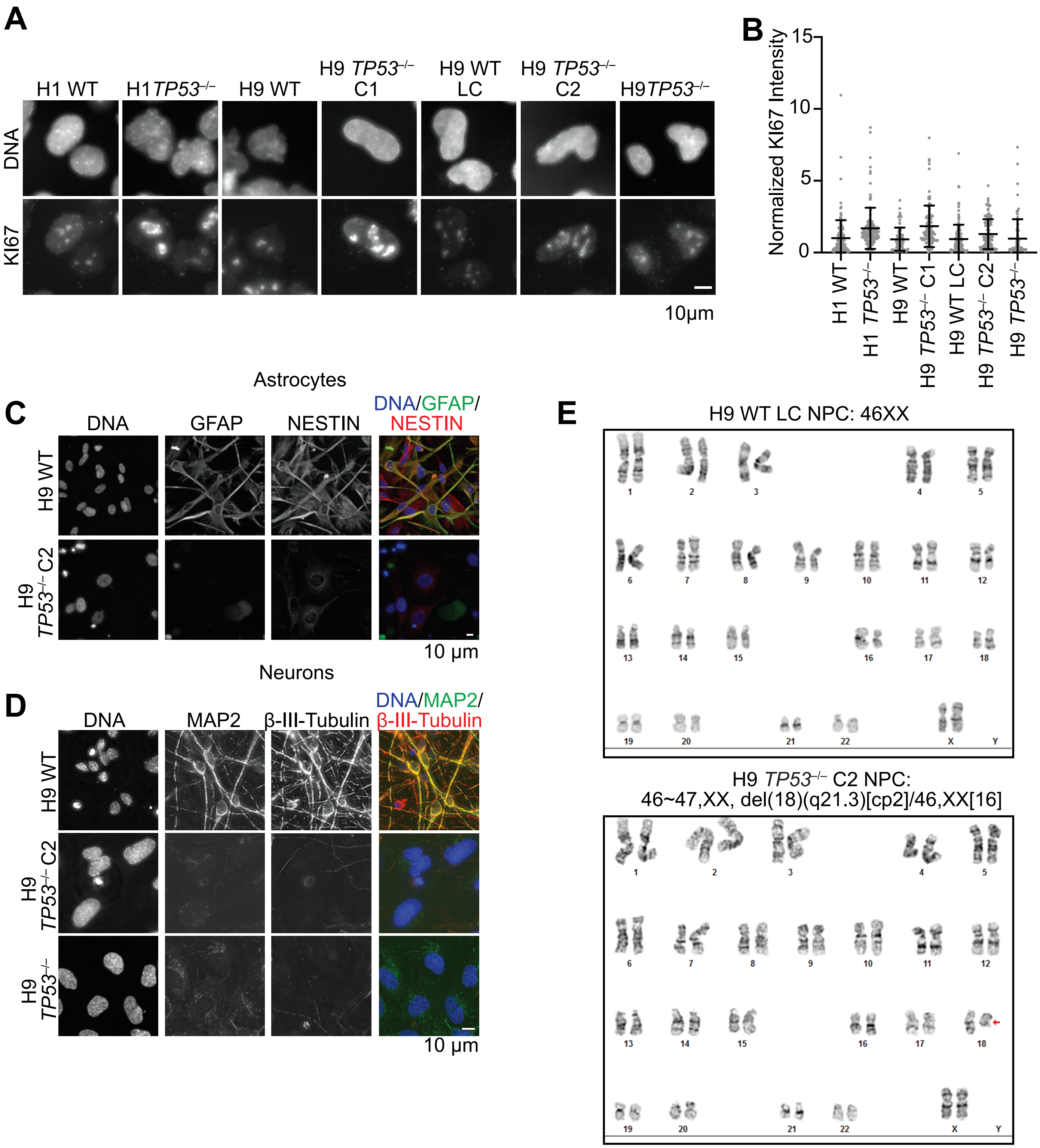

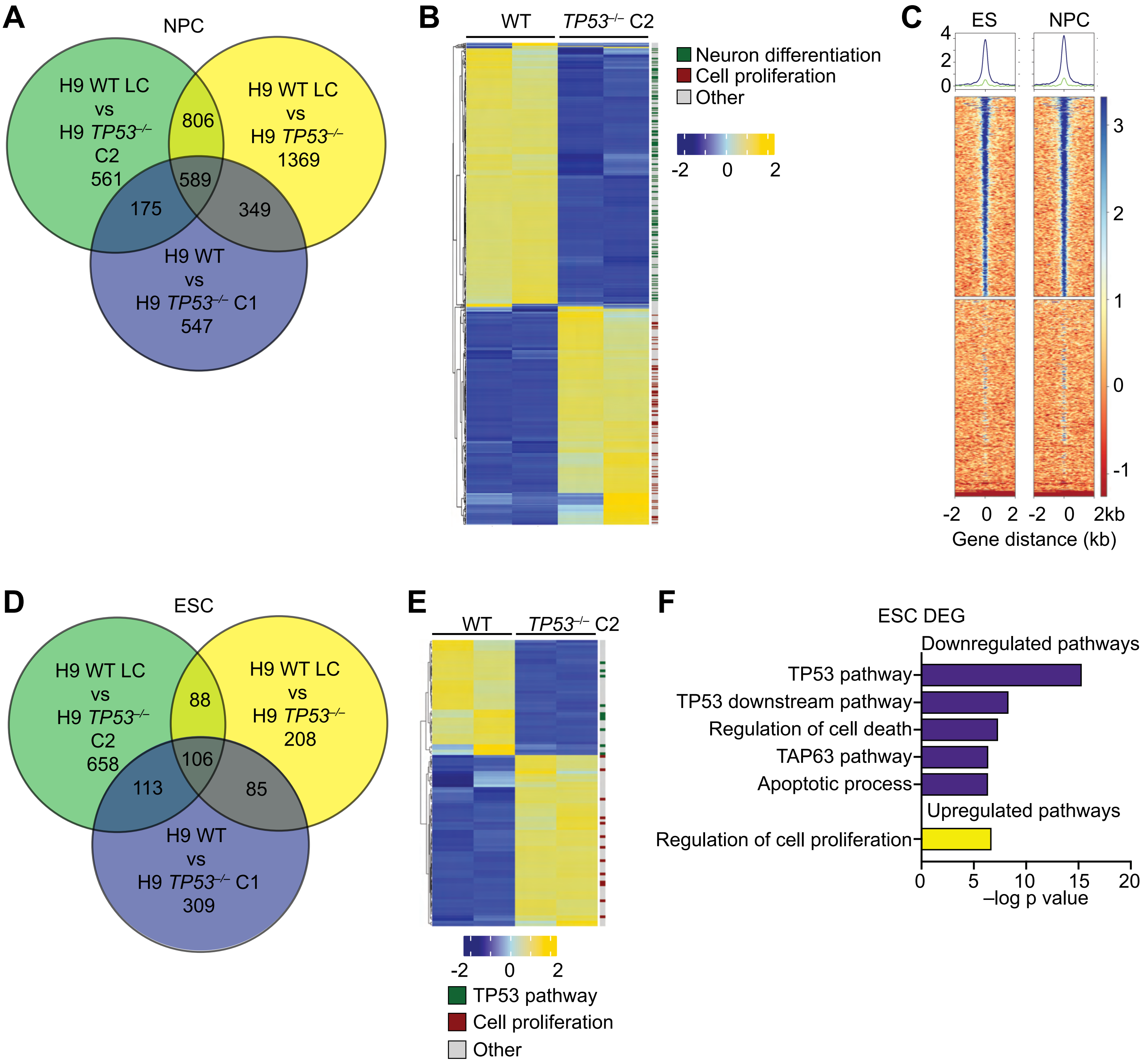

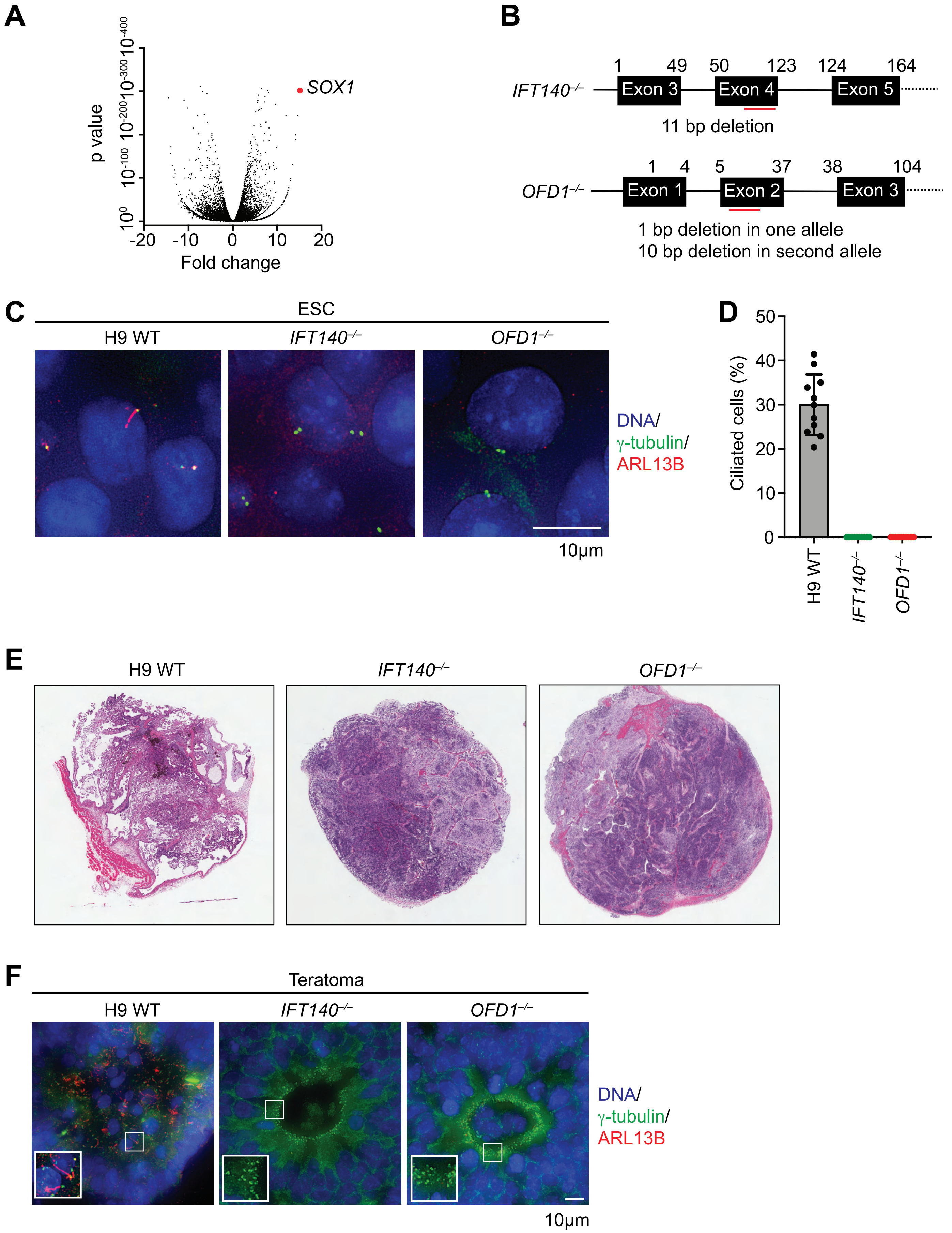

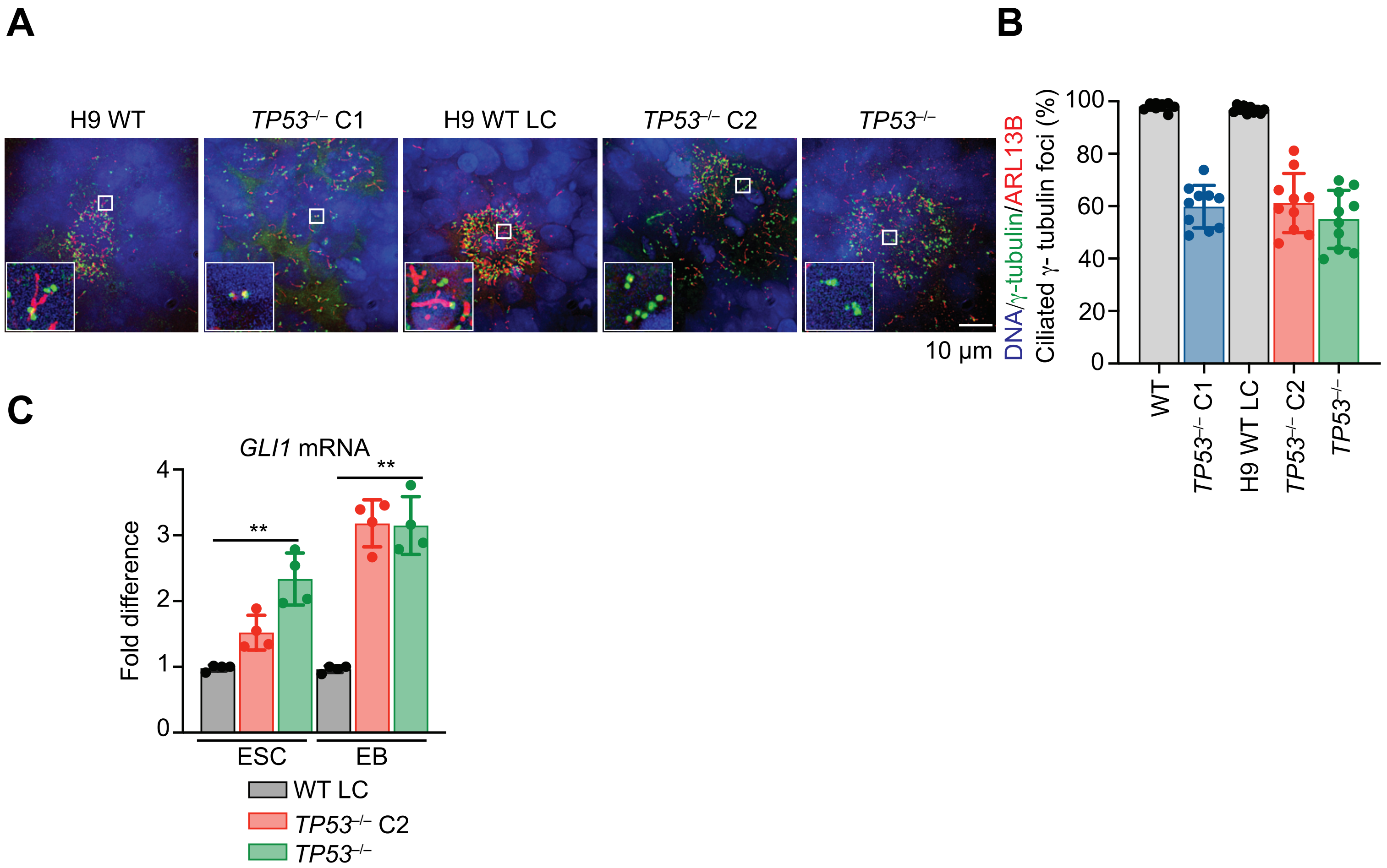

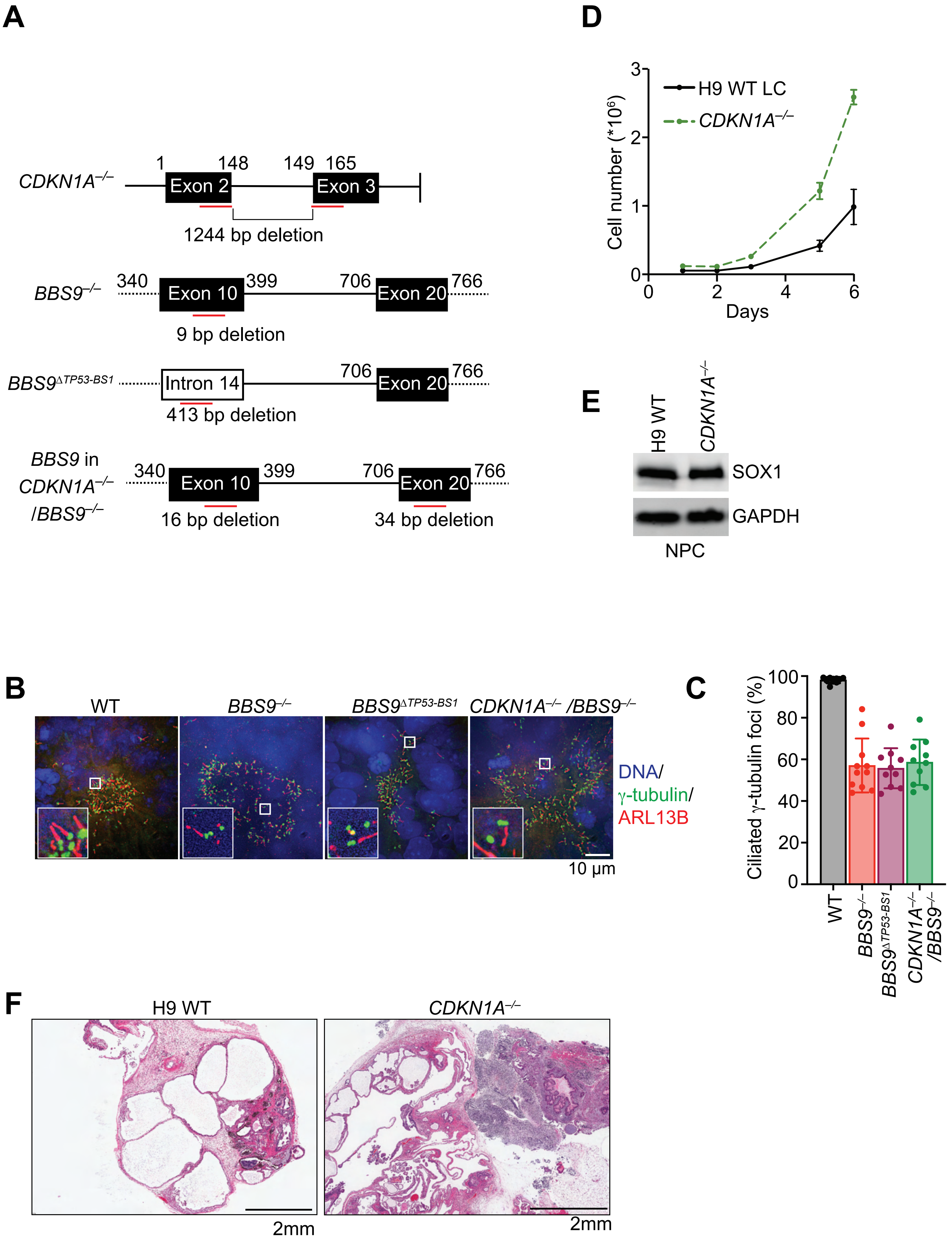
